# Astrocytic AMPK couples metabolic signals to hypothalamic circadian timing

**DOI:** 10.1101/2025.08.14.670385

**Authors:** María Luengo-Mateos, Antía González-Vila, María Silveira-Loureiro, Daniela Sofía Abreu, Mikel Azkargorta, Ali Mohammad Ibrahim-Alasoufi, Victor Pardo-García, Hugo López-Goldar, Paula Martínez-Pérez, Helena Covelo Molares, Nathalia R.V. Dragano, Paola Fernández- Sanmartín, Ánxela M Estévez-Salguero, Mariana Astiz, Cláudia Cavadas, Carlos Diéguez, Miguel Fidalgo, Diana Guallar, Félix Elortza, Joao Filipe Oliveira, Miguel López, Olga Barca-Mayo

## Abstract

Circadian clocks coordinate behaviour and physiology with daily cycles of light and nutrient availability, but how metabolic signals influence brain timing remains incompletely understood. Astrocytes integrate metabolic and hormonal cues and exhibit time-of-day-dependent responses, suggesting that they may convey temporal information to hypothalamic circuits. Here, we show that hypothalamic AMP-activated protein kinase (AMPK) exhibits circadian regulation independently of light and feeding cues, is modulated by nutrient availability and astrocytic Ca²⁺ signalling, and regulates the temporal organisation of the hypothalamic phosphoproteome. Genetic manipulation of astrocytic AMPK signalling alters PER2 abundance and phosphorylation, including at a conserved residue implicated in circadian period regulation. At the behavioural level, AMPK and PER2 in ventromedial hypothalamic astrocytes contribute to food-anticipatory activity, whereas disruption of astrocytic AMPK signalling alters SCN-dependent circadian locomotor rhythms and energy homeostasis in a sex-dependent manner. Together, these findings identify astrocytic AMPK signalling as a temporally regulated pathway that couples metabolic signals to hypothalamic circadian timing and systemic homeostasis.

## Introduction

Temporal coordination of behaviour and physiology is essential for maintaining organismal homeostasis^1^. In mammals, this coordination relies on a distributed network of cellular clocks present in nearly all tissues and hierarchically synchronized by the suprachiasmatic nucleus (SCN). Not all circadian rhythms, however, are light dependent. Scheduled feeding, for example, induces food-anticipatory activity (FAA), a surge in locomotion prior to mealtime that persists after SCN ablation, consistent with a food-entrainable timing mechanism^2^. Despite decades of behavioural evidence, the anatomical, cellular and molecular basis of FAA remains unresolved, and current models propose distributed food-sensitive timing networks across central and peripheral tissues^2^.

At the molecular level, circadian clocks rely on transcription-translation feedback loops in which positive (CLOCK, BMAL1) and negative (PER, CRY) regulators generate ∼24-h rhythms in gene expression^3^. Beyond transcription, temporal information can also be encoded through post-translational mechanisms, with phosphorylation emerging as a major and evolutionarily conserved regulatory layer^4^. In mammals, phosphoproteomic studies, conducted largely in peripheral tissues, suggest that phosphorylation rhythms can exceed those of transcripts or total proteins, identifying phosphoregulation as a prominent layer of temporal control^4–6^. However, how hypothalamic phosphosignalling is temporally organised, and how it integrates metabolic cues with circadian timing, remains largely unknown.

Astrocytes integrate hormonal, synaptic and metabolic cues^7–9^ and are increasingly recognized as active components of the circadian system. In the SCN, they exhibit intrinsic circadian oscillations and daily Ca²⁺ rhythms, modulate neuronal synchrony and contribute to behavioural and physiological rhythms^10–20^. Light is the dominant environmental signal that entrains SCN neurons through well-defined retinal and hypothalamic pathways^3^, yet the signals that entrain astrocyte clocks remain poorly understood. This gap is particularly relevant for food-entrainable rhythms, where metabolic and hormonal signals are thought to convey temporal information to the circadian system. However, the intracellular pathways that translate these metabolic inputs into circadian timing remain unresolved. AMP-activated protein kinase (AMPK) is well positioned to fulfil this role because it integrates metabolic and Ca²⁺ signals, has established roles in circadian and metabolic regulation, and is highly expressed in astrocytes^21–28^.

Here, we show that hypothalamic AMPK signalling exhibits circadian variation independently of light and feeding cues, is modulated by nutrient availability and Ca²⁺ signalling in astrocytes, and that astrocytic AMPK regulates the temporal organisation of the hypothalamic phosphoproteome. Perturbation of astrocytic AMPK signalling alters SCN-dependent circadian locomotor behaviour, PER2 abundance and phosphorylation, and metabolic homeostasis. In addition, AMPK signalling and PER2 in ventromedial hypothalamic (VMH) astrocytes regulate FAA. Together, these findings identify astrocytic AMPK signalling as a temporally regulated pathway that couples metabolic signals to hypothalamic circadian timing and systemic energy homeostasis.

## Results

### Astrocytic AMPKα1 regulates circadian locomotor behaviour

AMPK is a heterotrimeric complex composed of catalytic α and regulatory β and γ subunits^29^. To investigate whether astrocytic AMPK regulates circadian function, we first examined its distribution and the consequences of astrocytic AMPKα1 deletion on circadian behaviour. Phosphorylated acetyl-CoA carboxylase (pACCα), a canonical downstream target of AMPK, was enriched in the SCN core **(Fig. 1a)**, where vasoactive intestinal polypeptide (VIP)-expressing neurons have been implicated in network synchrony and circadian rhythm amplitude^29^. We generated an inducible Glast-CreERT2 conditional knockout of *Prkaa1* (hereafter referred to as *α1cKO*). Previous studies have shown that this approach targets astrocytes in the adult brain with minimal recombination in NeuN-positive neurons^9,15,16,19,30^. In *α1cKO* hypothalamus, AMPK signalling was reduced, with immunoblotting confirming decreases in AMPKα1, phosphorylated AMPKα (pAMPKα) and pACCα **(Fig. 1b and Supplementary Fig. 1a).**

**Fig. 1.**
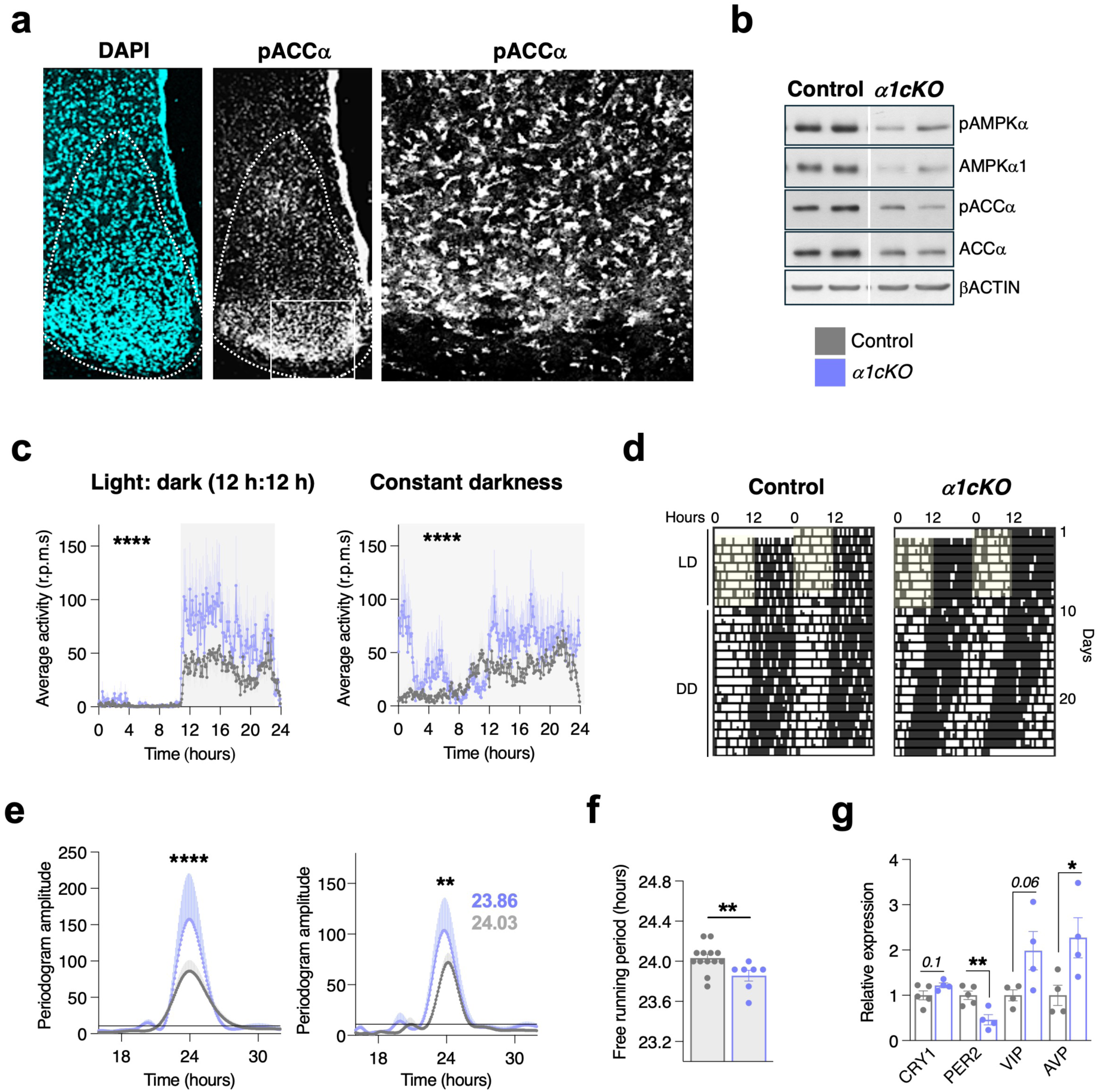
Astrocytic AMPKα1 loss regulates circadian locomotor behaviour. a, Representative immunofluorescence images showing pACCα in the SCN at ZT4 following overnight fasting. DAPI staining delineates SCN structure; the middle panel shows pACCα immunoreactivity and the right panel shows a higher-magnification view of the boxed ventrolateral SCN region. **b,** Representative western blots of AMPK signalling components in hypothalamic lysates from control and *α1cKO* mice collected at ZT8. **c,** Average locomotor activity profiles of control and *α1cKO* mice under 12 h:12 h light-dark conditions (LD) and constant darkness (DD) (control n = 13, *α1cKO* n = 7). **d,** Representative double-plotted actograms under LD and DD conditions. **e,** Lomb-Scargle periodogram analysis of locomotor activity rhythms (control n = 13, *α1cKO* n = 7), **f,** Free-running period under DD conditions (control n = 13, *α1cKO* n = 7). **g,** Quantification of CRY1, PER2, VIP and AVP protein abundance in hypothalamic tissue collected at ZT8 (n = 4-5). Data are presented as mean ± s.e.m. Statistical significance was determined using two-way ANOVA (c, e), and unpaired two-tailed t-tests (f, g). *P < 0.05, **P < 0.01, ****P < 0.0001.

Wheel-running rhythms were used as an index of SCN-driven circadian function^31,32^. To assess the impact of astrocytic AMPKα1 loss, we monitored wheel-running rhythms of *α1cKO* and floxed control mice under 12:12 h light-dark (LD) conditions followed by constant darkness (DD). Male *α1cKO* mice showed increased wheel-running activity, particularly under DD conditions, together with a shortening of the free-running period relative to matched floxed controls **(Fig. 1c-f and Supplementary Fig. 1b)**. At the molecular level, *α1cKO* hypothalamus exhibited elevated arginine vasopressin (AVP) and a trend toward increased VIP (p = 0.06), indicating altered SCN-associated neuropeptide signalling **(Fig. 1g and Supplementary Fig. 1c)**. Whereas in peripheral tissues AMPK promotes CRY1 and PER2 degradation^33–35^, in *α1cKO* hypothalamus PER2, but not CRY1, was selectively reduced, indicating that astrocytic AMPK signalling is required to maintain normal hypothalamic PER2 abundance. Astrocytic *Per2* deletion has also been reported to shorten free-running period^36^, suggesting that altered PER2 levels may contribute to some aspects of the behavioural phenotype observed in *α1cKO* mice. Together, these findings establish astrocytic AMPKα1 as a regulator of circadian locomotor behaviour and hypothalamic clock function.

### Astrocytic AMPKγ2 loss alters circadian locomotor behaviour and temporal organisation of the hypothalamic phosphoproteome

The regulatory AMPKγ2 subunit contributes to AMP sensing and AMPK activation dynamics, and activating mutations in PRKAG2 are associated with sustained AMPK activity^37–39^. Reanalysis of published circadian transcriptome data from CBA/CaJ mice^40^ revealed that *Prkag2* transcripts are rhythmic in the SCN, peaking at night, but not in other brain regions, identifying *Prkag2* as a candidate circadian regulator in the SCN **(Supplementary Fig. 2a)**. To test whether astrocytic AMPKγ2 deletion produces circadian phenotypes distinct from those observed after AMPKα1 loss, we generated an inducible Glast-CreERT2-based knockout of *Prkag2* (*γ2cKO*).

In control animals, hypothalamic pAMPKα levels were higher at ZT20 than at ZT8, in agreement with previous reports of nocturnal AMPK activation^34^. This temporal variation was blunted in *γ2cKO* animals, which showed comparable pAMPKα levels at both time points **(Supplementary Fig. 2b)**. In contrast, pACCα levels did not differ markedly between ZT8 and ZT20 in control hypothalami, whereas mutants exhibited elevated pACCα levels at ZT8 **(Supplementary Fig. 2b)**. In control hypothalami, pAMPKα and pACCα displayed distinct profiles across the two time points examined, consistent with previous observations in liver showing a temporal separation between AMPK activation and ACCα phosphorylation^5^. In *γ2cKO*;*tdTomato* reporter mice, pACCα immunoreactivity was increased in Tomato⁺ VMH astrocytes at ZT0, consistent with altered AMPK pathway activity in recombined cells **(Supplementary Fig. 2c)**. Expression of AMPKγ1, previously proposed to compensate for AMPKγ2 loss^41^, was reduced at ZT8 and unchanged at ZT20 **(Supplementary Fig. 2b)**, arguing against compensatory upregulation under these conditions. Together, these findings demonstrate disrupted temporal organisation of hypothalamic AMPK signalling in *γ2cKO* mice.

To assess the impact of astrocytic AMPKγ2 loss on circadian locomotor activity, we monitored wheel-running rhythms of *γ2cKO* and floxed control mice. Under LD conditions, *γ2cKO* mice exhibited reduced nocturnal locomotor activity and lower rhythm amplitude, whereas total daily locomotor activity remained largely preserved **(Fig. 2a, b and Supplementary Fig. 2d)**, indicating that mutants do not exhibit a generalized hypoactivity phenotype. Under DD conditions, *γ2cKO* animals displayed a significant lengthening of the free-running period relative to matched γ2 floxed controls (23.83 ± 0.06 h versus 24.08 ± 0.02 h; P = 0.001; **Fig. 2a, b and Supplementary Fig. 2d**). Because free-running periods differed modestly between independently generated α1 and γ2 floxed cohorts, we additionally performed a pooled analysis across floxed controls, which confirmed a moderate but significant period lengthening in *γ2cKO* mice (P < 0.05). Relative to pooled controls (23.96 ± 0.04 h), *α1cKO* mice showed a modest shortening of free-running period (23.86 ± 0.05 h), which did not reach statistical significance (adjusted P = 0.18). Together, these findings demonstrate that altered astrocytic AMPK signalling regulates circadian locomotor timing, with astrocytic AMPKγ2 loss producing robust changes in free-running period.

**Fig. 2.**
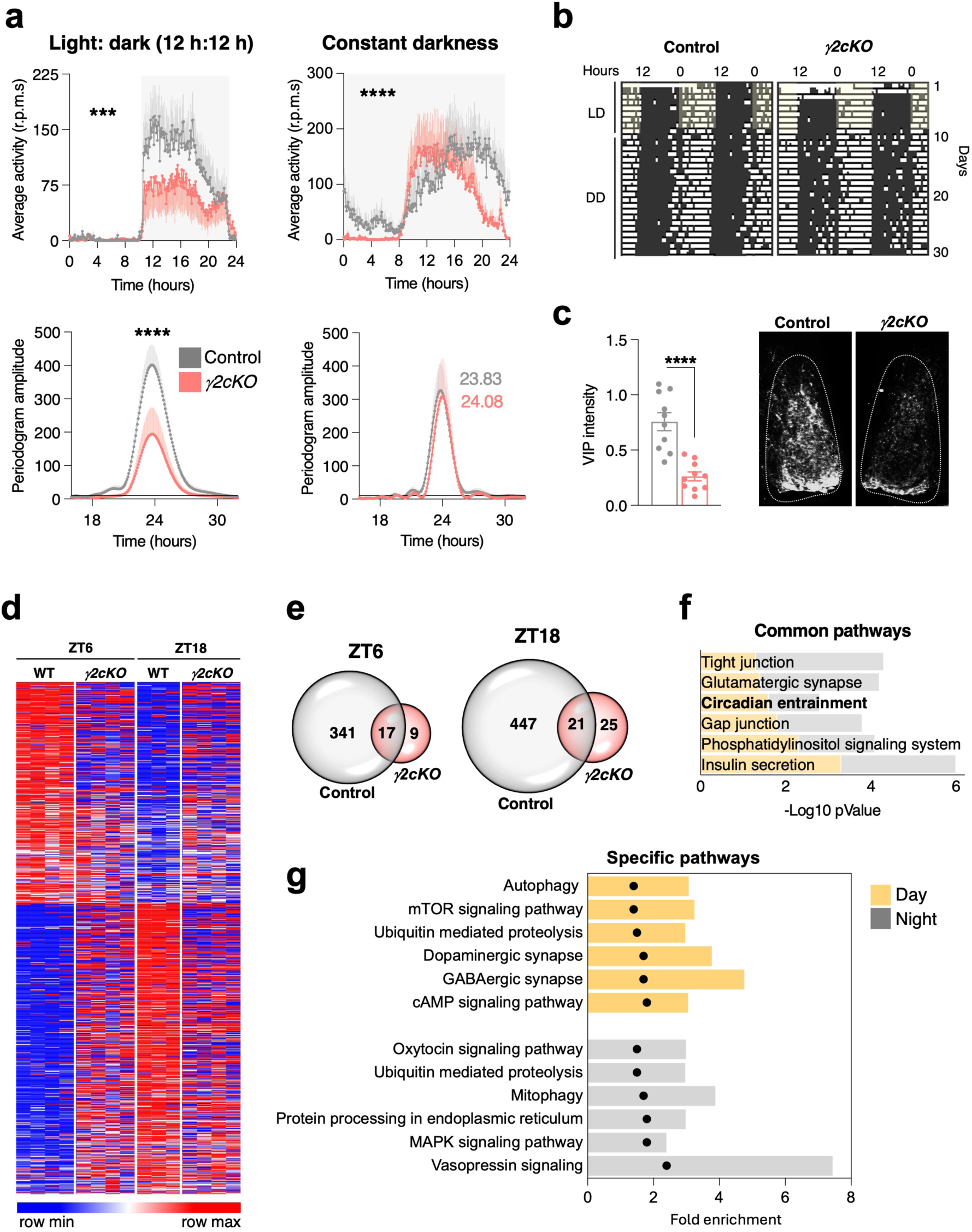
Astrocytic AMPKγ2 loss alters circadian locomotor behaviour and temporal organisation of the hypothalamic phosphoproteome. a, Average locomotor activity profiles of control and *γ2cKO* mice under 12 h:12 h LD and DD (upper panels). Lomb-Scargle periodogram analysis of locomotor activity rhythms under LD and DD (lower panels) (n = 8). **b,** Representative double-plotted actograms from control and *γ2cKO* mice under LD and DD conditions. **c,** Quantification of VIP immunoreactivity in the SCN at ZT2 (left) and representative confocal images from control and *γ2cKO* mice (right) (n = 10). **d,** Heatmap showing phosphoproteins displaying significant time-of-day-dependent variation between ZT6 and ZT18 in control and *γ2cKO* hypothalami (control n = 3; *γ2cKO* n = 4). **e,** Venn diagrams showing the number of phosphoproteins displaying significant time-of-day-dependent variation between ZT6 and ZT18 in control and *γ2cKO* hypothalami. **f,** KEGG pathway enrichment analysis of pathways associated with phosphoproteins showing time-of-day-dependent variation in control hypothalami but not in *γ2cKO* mice. Pathways enriched at both ZT6 and ZT18 are shown. **g,** KEGG pathway enrichment analysis of phase-specific pathways associated with phosphoproteins showing time-of-day-dependent variation in control hypothalami but not in *γ2cKO* mice. Yellow and grey indicate pathways preferentially enriched at ZT6 and ZT18, respectively. Bars indicate fold enrichment and black dots indicate −log_10_(*P* value). Data are presented as mean ± s.e.m. Statistical significance was determined using two-way ANOVA (a) and unpaired two-tailed t-tests (c). \*\*\**P* < 0.001, \*\*\*\**P* < 0.0001.

Notably, *γ2cKO* mice showed reduced SCN VIP immunoreactivity **(Fig. 2c)**. To further assess the contribution of SCN astrocytic AMPKγ2 signalling to circadian behaviour, we examined locomotor rhythmicity after virogenetic, SCN-restricted astrocytic recombination, anatomically validated by GFP staining **(Supplementary Fig. 3a)**. This manipulation markedly reduced locomotor rhythm amplitude under both LD and DD conditions and produced a significant lengthening of the free-running period **(Supplementary Fig. 3b-d)**. Despite the attenuation of rhythmic amplitude, residual locomotor activity persisted across the circadian cycle, indicating that disruption of astrocytic AMPKγ2 within the SCN weakens behavioural circadian rhythms. Together, these findings establish SCN astrocytic AMPKγ2 as a regulator of circadian locomotor behaviour.

Having established that astrocytic AMPKγ2 loss disrupts hypothalamic AMPK pathway dynamics and circadian locomotor behaviour, we next asked how this manipulation affects the temporal organisation of the hypothalamic phosphoproteome. In control animals, phosphorylation profiles differed substantially between ZT6 and ZT18, with more than 350 phosphoproteins showing significant phase-associated variation. In *γ2cKO* hypothalami, these phase-dependent phosphorylation differences were largely abolished **(Fig. 2d, e)**, demonstrating a global disruption of temporal phosphoregulation. KEGG pathway analyses revealed that phosphoproteins displaying temporal regulation in controls but disrupted rhythmicity in *γ2cKO* mice converged on pathways involved in circadian entrainment, synaptic signalling, phosphatidylinositol signalling and insulin secretion **(Fig. 2f)**. Additional pathways displaying phase-specific temporal phosphoregulation included metabolic regulation (mTOR signalling, autophagy and mitophagy), neurotransmission (dopaminergic and GABAergic synapses and cAMP signalling) and neuropeptide signalling (vasopressin and oxytocin pathways) **(Fig. 2g)**. Independent comparisons between control and *γ2cKO* mice at each circadian phase revealed widespread genotype-dependent phosphoproteomic alterations **(Supplementary Fig. 3e-g)**. At ZT6, differentially regulated phosphoproteins were enriched in pathways related to neurotransmitter and ion transport, biological rhythms and apoptosis, whereas at ZT18 they were associated with RNA processing, intracellular trafficking and cell cycle-related functions. Thus, beyond disrupting the temporal phosphorylation programmes normally observed in control hypothalami, astrocytic AMPKγ2 deficiency produces distinct phase-dependent signalling alterations across circadian time. Collectively, these findings establish astrocytic AMPKγ2 as a regulator of the temporal organisation of the hypothalamic phosphoproteome and circadian locomotor behaviour.

### Astrocytic AMPK regulates PER2 temporal dynamics

Building on the observation that astrocytic AMPKα1 loss reduces hypothalamic PER2 **(Fig. 1g and Supplementary Fig. 1c)**, we next examined PER2 abundance in *γ2cKO* hypothalami at ZT8 and ZT20. In control mice, total PER2 was higher at ZT20 than at ZT8, whereas in *γ2cKO* mice PER2 levels were already elevated at ZT8, demonstrating altered temporal regulation of PER2 abundance in the absence of astrocytic AMPKγ2 **(Fig. 3a)**.

**Fig. 3.**
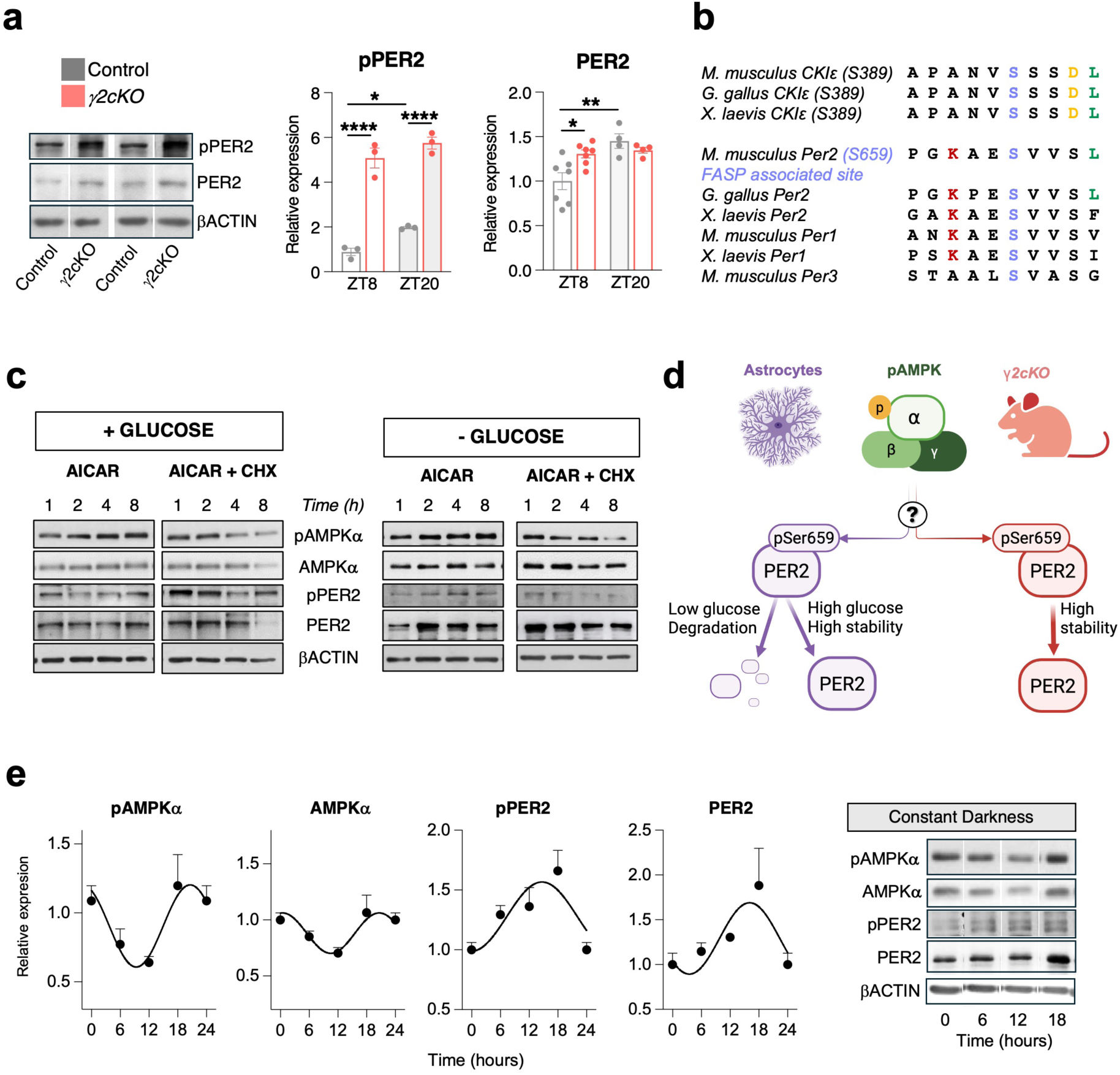
Astrocytic AMPK regulates PER2 temporal dynamics. a, Representative immunoblots (left) and quantification (right) of pPER2 and total PER2 protein levels in hypothalamic lysates from male control and *γ2cKO* mice collected at ZT8 and ZT20 (n = 3-7). **b,** Sequence alignment of PER2 Ser659 and surrounding residues across vertebrates, highlighting conservation of the AMPK consensus motif and CKIδ/ε target sites. **c,** Representative immunoblots of primary hypothalamic astrocytes treated with the AMPK activator AICAR under high-glucose (+ glucose) or low-glucose (− glucose) conditions, with or without cycloheximide (CHX), for the indicated times (1, 2, 4 and 8 h). Representative of two independent experiments. **d,** Schematic model summarizing the relationship between astroglial AMPK activity and PER2 dynamics under different metabolic conditions. **e,** Circadian expression profiles of pAMPKα, total AMPKα, pPER2 and total PER2 in hypothalamic lysates collected from control mice under constant darkness. Representative immunoblots are shown (right). Peak phases derived from cosine fitting are indicated in each plot (n = 4-6). Data are presented as mean ± s.e.m. Statistical significance was determined using two-way ANOVA (a). *P < 0.05, **P < 0.01, ****P < 0.0001.

To place these changes in a mechanistic context, we focused on Ser659, a conserved residue within the familial advanced sleep phase (FASP) phosphoregulatory region of PER2, homologous to human Ser662. This site is implicated in CK1-dependent control of PER2 stability and circadian period^42^. The surrounding serine-rich region is targeted by CK1ε and contains a sequence compatible with the AMPK consensus motif [ϕ-X-(R/K)-X-X-S/T]^27^ **(Fig. 3b).** We therefore examined Ser659-phosphorylated PER2 (pPER2) in control and *γ2cKO* hypothalami. In control mice, pPER2 levels were higher at ZT20 than at ZT8, whereas *γ2cKO* mice exhibited elevated pPER2 levels already at ZT8, mirroring the changes observed for total PER2 and demonstrating altered temporal regulation of PER2 phosphorylation.

To probe how AMPK pathway activation and metabolic state influence PER2 dynamics in astrocytes, we treated primary hypothalamic astrocytes with the AMPK activator AICAR under high-or low-glucose conditions **(Fig. 3c)**. Low glucose lowers cellular energy charge and thereby promotes AMPK activation, whereas AICAR provides a pharmacological stimulus to the pathway. AICAR robustly increased pAMPK under both glucose conditions. Under high glucose, pPER2 transiently decreased at 2-4 h and rebounded by 8 h, whereas total PER2 remained relatively stable. Under low glucose, pPER2 and PER2 paralleled pAMPK dynamics more closely, rising over time and declining at 8 h. To assess whether sustained PER2 levels require ongoing protein synthesis, we performed the same experiment in the presence of cycloheximide (CHX). Under these conditions, pAMPK, pPER2 and total PER2 all declined after AICAR treatment, indicating that sustained PER2 levels over these timescales require ongoing protein synthesis and are not maintained by AMPK pathway activation alone. Although these data do not distinguish direct PER2 phosphorylation from an indirect mechanism, they demonstrate that PER2 dynamics depend on both AMPK pathway activation and glucose availability. Previous studies have likewise shown that glucose availability can alter PER2 rhythmicity in neurons, highlighting the broader sensitivity of clock protein dynamics to metabolic state^43^ **(Fig. 3d).**

To assess whether similar temporal relationships were also evident *in vivo*, we examined PER2 and pPER2 in hypothalamic tissue collected under DD conditions. Both total PER2 and pPER2 reached their highest levels around ZT20, recapitulating the temporal pattern observed under LD conditions and coinciding with elevated pAMPK levels **(Fig. 3e)**. Taken together, these findings identify astrocytic AMPK signalling as a regulator of PER2 temporal dynamics and provide a potential link between metabolic state and hypothalamic circadian timing.

### Astrocytic AMPKγ2 regulates sex-specific metabolic homeostasis

Altered AMPKγ2 signalling has previously been associated with obesity, hyperphagia and systemic metabolic dysfunction in both humans and animal models^37–39,44^. However, whether astrocyte-specific loss of AMPKγ2 signalling is sufficient to impair systemic metabolic coordination remained unknown. Because sex-dependent differences have been reported in metabolic and circadian homeostasis^9,19,45–48^, we evaluated the impact of astrocytic AMPKγ2 deletion in female mice. Female *γ2cKO* mice showed preserved free-running period, nocturnal activity and total locomotion **(Supplementary Fig. 4a, b)**. Nevertheless, pAMPK and pACCα failed to exhibit normal day-night variation in *γ2cKO* female hypothalami **(Supplementary Fig. 4c)**, indicating disrupted temporal AMPK regulation in the absence of overt behavioural circadian alterations. Notably, female *γ2cKO* mice exhibited progressive weight gain accompanied by increased fat and lean mass, without detectable alterations in food intake, locomotor activity or respiratory quotient **(Fig. 4a and Supplementary Fig. 5a-c)**. In addition, *γ2cKO* female mice exhibited reduced energy expenditure and impaired thermogenesis, evidenced by lower rectal and brown adipose tissue (BAT) temperatures together with reduced BAT uncoupling protein 1 (UCP1) expression **(Fig. 4b-d and Supplementary Fig. 5d, e)**. Fasting glucose remained normal, although glucose and insulin tolerance tests revealed modest impairments in glycaemic control **(Fig. 4e, f and Supplementary Fig. 5f)**. Thus, disruption of astroglial AMPK signalling impairs energy balance in females despite preserved locomotor circadian outputs. Of note, male *γ2cKO* mice developed a more severe metabolic phenotype, characterised by weight gain, increased adiposity, hyperphagia, reduced dark-phase activity, decreased energy expenditure, elevated respiratory quotient and marked insulin resistance **(Fig. 4 g-k and Supplementary Fig. 5 h-j)**. Notably, survival was reduced in both sexes, but more markedly in males **(Fig. 4l and Supplementary Fig. 5g)**, revealing a marked male vulnerability to chronic disruption of astroglial AMPK signalling. Together, these findings establish astroglial AMPK signalling as a sex-dependent regulator of systemic metabolic homeostasis.

**Fig. 4.**
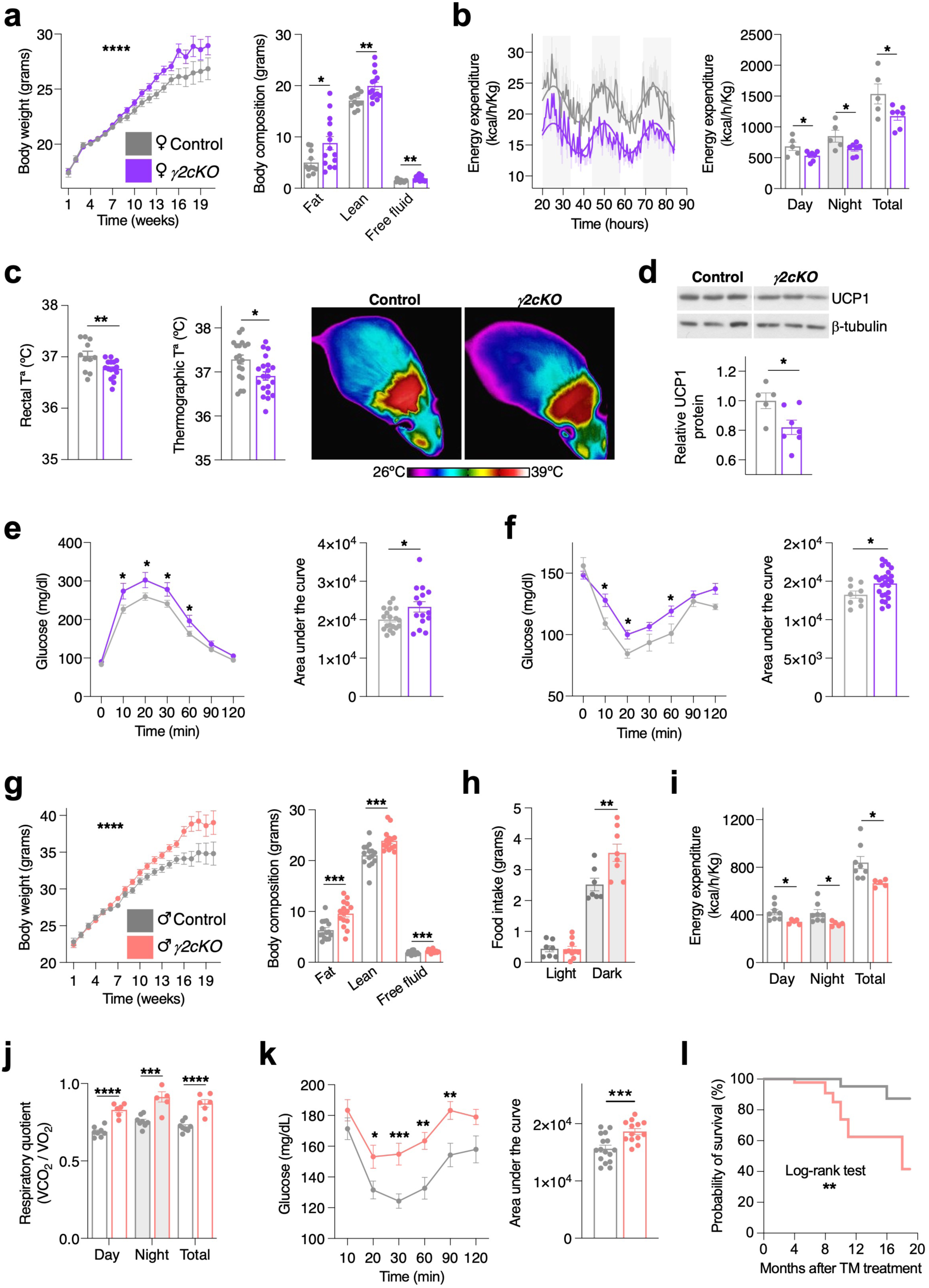
Astrocytic AMPKγ2 regulates sex-specific metabolic homeostasis. **a**, **g**, Body weight trajectories and body composition analysis in control and *γ2cKO* females (**a**) and males (**g**) (females n = 11-18; males n = 15-18). **b,** Energy expenditure profiles over 72 h (left) and quantification of average day-time, night-time and total energy expenditure (right) in females (n = 5-7). **c,** Rectal temperature (left), BAT surface temperature quantified by infrared thermography (middle) and representative infrared images (right) in females (n = 11-20). **d,** Representative immunoblots (top) and quantification (bottom) of UCP1 protein abundance in BAT from females (n = 5-7). **e, f,** Glucose tolerance test (**e**) and insulin tolerance test (**f**) with corresponding area under the curve (AUC) in females (GTT n = 15-19; ITT n = 9-25). **h,** Food intake during light and dark phases in males (n = 7-8). **i,** Quantification of average day-time, night-time and total energy expenditure in males (n = 5-8). **j,** Quantification of average day-time, night-time and total respiratory exchange ratio (RER) in males (n = 5-8). **k,** Insulin tolerance test and corresponding AUC in males (n = 13-16). **l,** Survival analysis of control and *γ2cKO* males (n = 94). Panels a-k were performed 20 weeks after tamoxifen induction, except for body weight trajectories. Data are presented as mean ± s.e.m. Statistical significance was determined using two-way ANOVA, unpaired two-tailed t-tests and the log-rank test. *P < 0.05, **P < 0.01, ***P < 0.001, ****P < 0.0001.

### VMH astrocytic AMPK and PER2 contribute to FAA

Regularly scheduled feeding elicits FAA, a preprandial increase in locomotion that persists after SCN ablation^2^. Given that astrocytes integrate metabolic cues and AMPK is a key cellular energy sensor, we hypothesised that astroglial AMPK signalling could contribute to FAA. To test this, control, *γ2cKO* and *α1cKO* mice were subjected to restricted feeding during the light phase (ZT4-10 for six days, followed by ZT4-8 for four days^9,19,45^) **(Fig. 5a).**

**Fig. 5.**
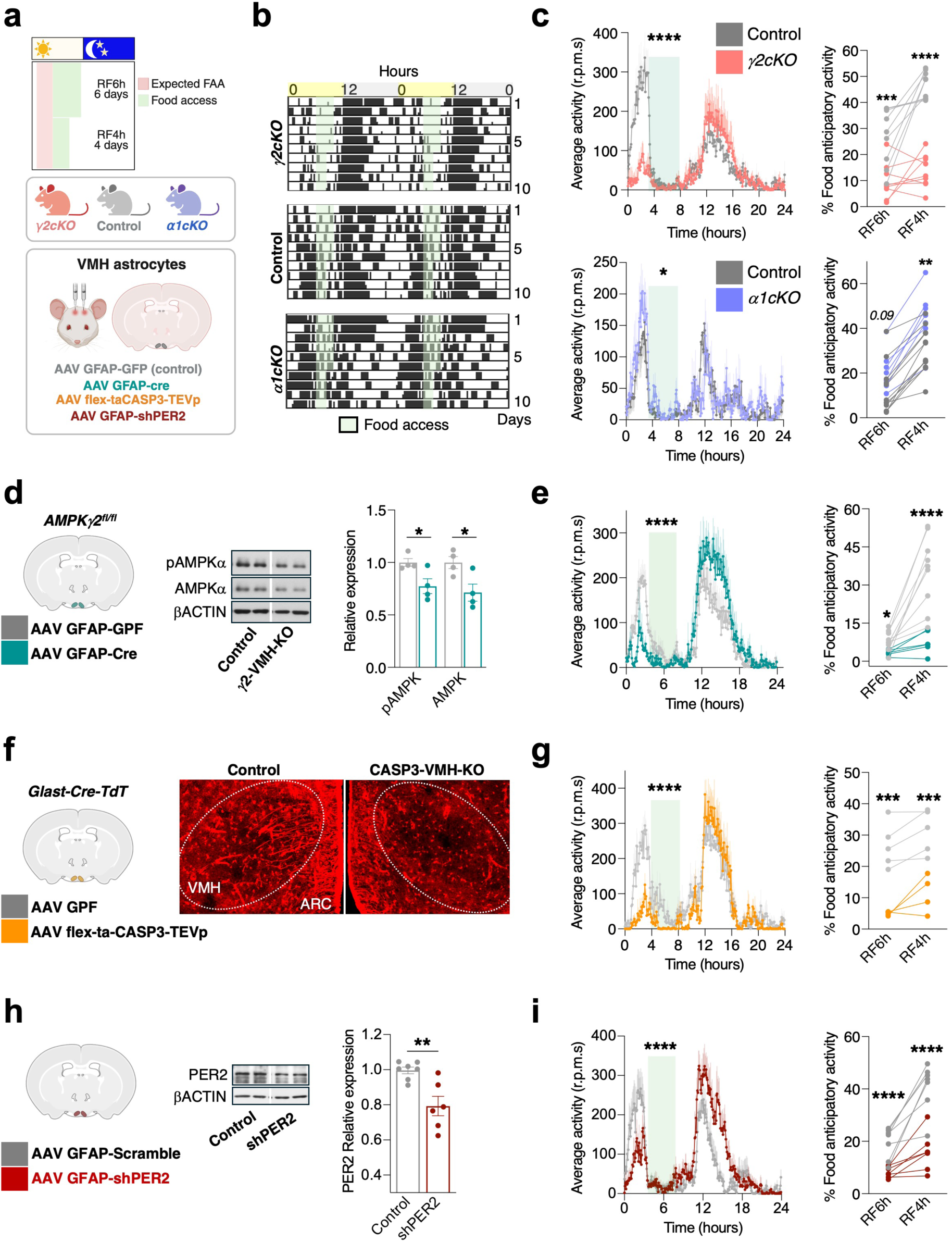
VMH astrocytic AMPK and PER2 contribute to FAA. a, Schematic of the restricted-feeding (RF) paradigm and experimental design for VMH-targeted viral manipulations. **b,** Representative actograms of locomotor activity during RF in *γ2cKO*, control and *α1cKO* mice. **c,** Average locomotor activity profiles (left) and FAA quantification (right) in *γ2cKO* (upper panel, n = 9) and *α1cKO* mice (lower panel, n = 6-12) during RF. **d,** Representative immunoblot (left) and quantification of pAMPKα and AMPKα (right) in VMH from *AMPKγ2^fl/fl^* mice injected with either AAV-GFAP-GFP or AAV GFAP-Cre (n = 4). **e,** Average locomotor activity profiles (left) and FAA quantification (right) following VMH-specific deletion of AMPKγ2 (n = 6-9). **f,** Representative VMH sections from *Glast^Cre-TdTomato^* mice showing astrocyte ablation after AAV-flex-taCASP3-TEVp injection. **g,** Average locomotor activity profiles (left) and FAA quantification (right) following VMH astrocyte ablation (n = 4-5). **h,** PER2 knockdown in VMH astrocytes using AAV-GFAP-shPER2. Representative immunoblot (left) and quantification of PER2 protein levels (right) (n = 6-7). **i,** Average locomotor activity profiles (left) and FAA quantification (right) in AAV-GFAP-shPER2-injected mice (n = 7). All locomotor activity profiles shown in panels c, e, g and i correspond to the RF4h phase following adaptation to RF6h. Data are presented as mean ± s.e.m. Statistical significance was determined using two-way ANOVA and unpaired two-tailed t-tests. *P < 0.05, **P < 0.01, ***P < 0.001, ****P < 0.0001.

As expected, control animals developed robust FAA, characterised by increased locomotor activity preceding food availability. In *γ2cKO* males, FAA was markedly reduced despite preserved total locomotor activity, whereas *α1cKO* mice displayed enhanced FAA accompanied by reduced nocturnal activity **(Fig. 5b, c and Supplementary Fig. 6a, b, d, e)**. These behavioural alterations were accompanied by distinct metabolic responses to restricted feeding: *γ2cKO* males retained higher body weight despite reduced food intake, whereas *α1cKO* mice were hyperphagic and lost substantial weight under restricted feeding conditions **(Supplementary Fig. 6c, f)**. Together, these bidirectional phenotypes demonstrate that temporally coordinated astroglial AMPK signalling is required for normal FAA expression. Despite progressive weight loss during restricted feeding, *γ2cKO* females retained higher body weight than controls and preserved FAA and behavioural rhythmicity, suggesting relative preservation of behavioural adaptation despite altered metabolic conditions^9,15,16,19,45^ **(Supplementary Fig. 6g-j)**.

At the molecular level, hypothalamic AMPK and ACCα phosphorylation in control animals were elevated at ZT4 relative to ZT6, consistent with phase-appropriate nutrient-responsive signalling. This temporal adjustment was blunted in *γ2cKO* males, which maintained elevated AMPK phosphorylation postprandially **(Supplementary Fig. 7a)**, indicating impaired adaptation of hypothalamic AMPK signalling to feeding-related temporal cues. Taken together, these findings demonstrate that hypothalamic AMPK signalling fails to undergo normal feeding-dependent temporal regulation in *γ2cKO* mice.

Previous studies have implicated the VMH in food anticipation and metabolic adaptation to restricted feeding ^49–53^. We therefore examined whether VMH astrocytes contribute directly to FAA regulation. Targeted deletion of AMPKγ2 in VMH astrocytes (AAV-GFAP-Cre in *Prkag2^fl^*^/*fl*^ mice) reduced local pAMPK and markedly reduced FAA, recapitulating the whole-brain *γ2cKO* phenotype **(Fig. 5d, e and Supplementary Fig. 7b)**. Similarly, selective astrocyte ablation within the VMH (AAV-flex-taCASP3-TEVp in *GlastCre*-*TdTomato* mice) attenuated FAA without substantially altering baseline locomotor behaviour **(Fig. 5f, g and Supplementary Fig. 7c).** Finally, astrocyte-specific knockdown of *Per2* within the VMH (AAV-GFAP-shPER2) also blunted FAA **(Fig. 5h, i and Supplementary Fig. 7d)**, indicating that PER2 in VMH astrocytes also contributes to FAA. This is consistent with previous evidence showing that global, but not neuron-specific, *Per2* deletion disrupts food anticipation^54,55^. Together, these findings identify AMPK signalling and PER2 in VMH astrocytes as critical regulators required for normal FAA expression.

### Astrocytic Ca2+ signalling contributes to temporal hypothalamic AMPK regulation

Although AMPK is canonically described as an energy sensor activated by elevated AMP/ATP ratios^23^, previous work^34^, together with our data **(Supplementary Fig. 2b)**, has shown that hypothalamic AMPK activity varies across the circadian cycle, peaking in the active phase despite largely stable brain ATP levels^56^. This pattern is consistent with AMPK being modulated by circadian mechanisms in addition to acute energetic state. To examine whether temporal changes in astrocytic AMPK phosphorylation can be observed in isolation, we synchronised primary hypothalamic astrocytes with insulin under low-or high-glucose conditions^9,16–18^ or treated them with AICAR **(Supplementary Fig. 8a)**. Insulin induced rhythmic AMPKα and ACCα phosphorylation under both glucose conditions and after AICAR treatment, and the overall phosphorylation profiles were influenced by glucose availability and AICAR exposure, suggesting that astrocytic AMPK phosphorylation rhythms retain intrinsic temporal regulation while remaining sensitive to metabolic state.

To separate nutrient-driven from circadian influences *in vivo*, we performed time-resolved hypothalamic phosphoproteomics in mice fasted for 24 h, with sampling across the circadian cycle. Under these conditions, 347 phosphoproteins showed time-of-day-dependent oscillations **(Fig. 6a, b)**, demonstrating rhythmic phosphorylation patterns even in the absence of feeding cues. Unsupervised clustering identified two phosphorylation peaks at ZT0 and ZT13, together with a prominent nocturnal module that was absent in *ad libitum*-fed animals. Kinase enrichment analysis of this night-time module highlighted AMPKα1 and AMPKα2 (*Prkaa1* and *Prkaa2*) among the top predicted upstream regulators **(Fig. 6c)**, suggesting that nutrient deprivation enhances AMPK-associated signalling at night.

**Fig. 6.**
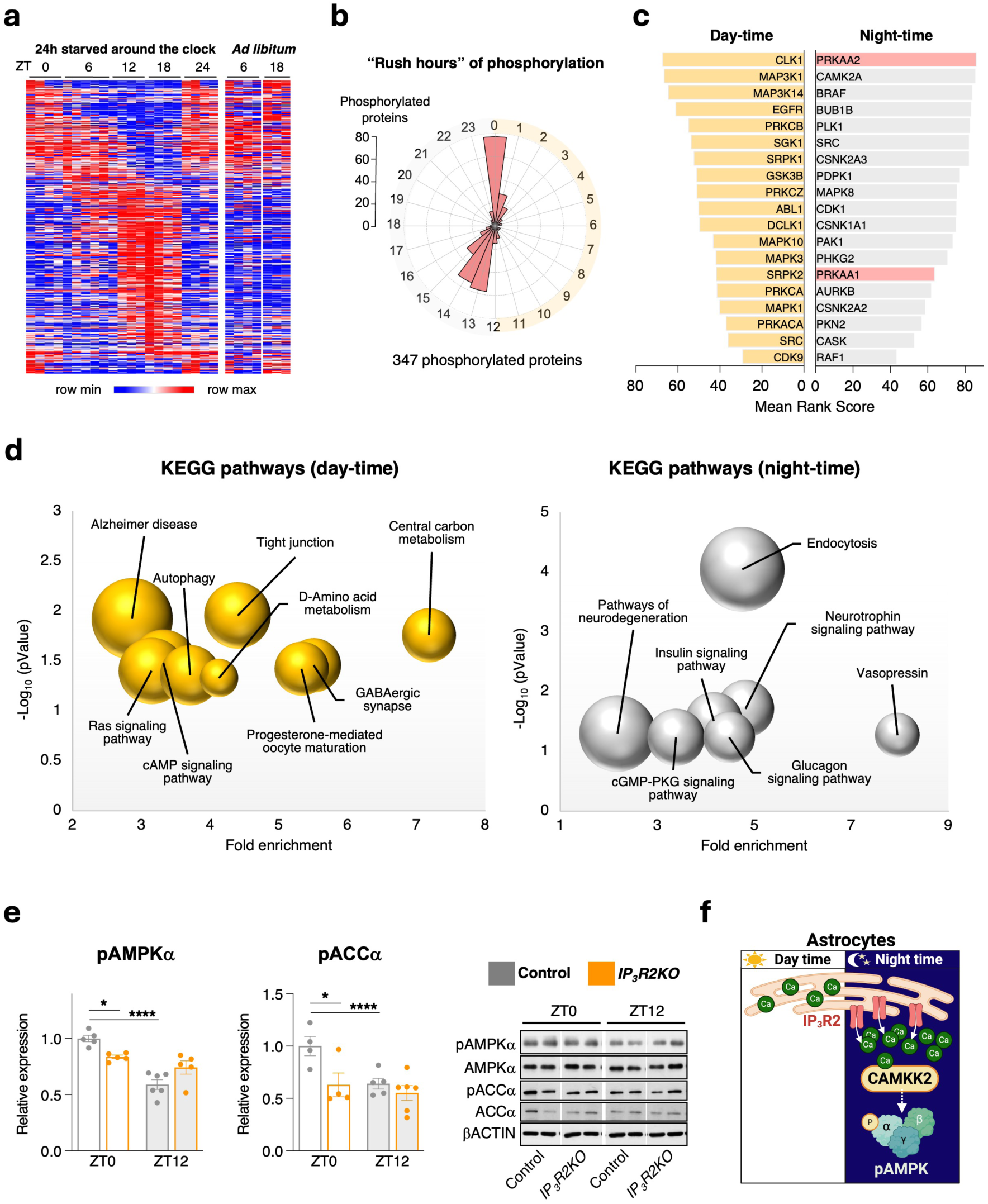
Astrocytic Ca^2+^ signalling contributes to temporal hypothalamic AMPK regulation. a, Heatmap of time-of-day-dependent hypothalamic phosphoproteins after 24-h fasting (left) or *ad libitum* feeding (right) across four circadian time points (n = 3-4 per group). **b,** Polar plot showing the phase distribution of the 347 rhythmic phosphoproteins detected under fasting conditions, with peaks in phosphorylation frequency at ZT0 and ZT13. **c,** Kinase-substrate enrichment analysis showing the top predicted upstream kinases for day-time (yellow) and night-time (grey) phosphorylation profiles; red text indicates AMPK subunits. **d,** KEGG pathway enrichment analysis of phosphoproteins enriched during the day (ZT0, yellow) or night (ZT13, grey). Bubble position indicates fold enrichment and statistical significance (−log_10_(P value)), whereas bubble size reflects the number of proteins assigned to each pathway. **e,** Quantification (left) and representative immunoblots (right) of pAMPKα and pACCα in hypothalamic extracts from control and *IP*₃*R2KO* mice under *ad libitum* feeding at ZT0 and ZT12 (n = 4-6 per group). **f,** Schematic model illustrating a potential role for astrocytic Ca^2+^-dependent signalling in the temporal regulation of AMPK activity. Night-time IP_3_R2-mediated Ca^2+^ release may promote CaMKK2-associated AMPK phosphorylation, whereas lower day-time Ca^2+^signalling is associated with reduced AMPK phosphorylation. Data are presented as mean ± s.e.m. Statistical significance was determined using two-way ANOVA. *P < 0.05, ****P < 0.0001.

Pathway enrichment analysis revealed a marked temporal segregation of phosphoproteins under fasting **(Fig. 6d)**. Day-time phosphoproteins mapped to GABAergic synapses and tight junctions, alongside pathways related to autophagy and cAMP signalling. In contrast, the night-time module was enriched for neuroendocrine and metabolic coordination pathways, including neurotrophin, glucagon, vasopressin and insulin signalling, suggesting a shift in functional priorities across the day-night cycle. Gene ontology analysis reinforced this partitioning: day-time phosphoproteins were dominated by glycolysis, translational regulation and solute symport, whereas night-time targets were enriched for transport processes, mRNA and rRNA processing, and ribosome biogenesis **(Supplementary Fig. 8b)**. Together, these data demonstrate that fasting uncovers a time-of-day-dependent organisation of hypothalamic phosphoregulation and suggest that nutrient deprivation reshapes AMPK-associated signalling networks across the circadian cycle, rather than AMPK acting solely as an acute energy sensor.

Ca^2+^/calmodulin-dependent protein kinase kinase 2 (CaMKK2) is highly expressed in hypothalamic circuits controlling energy balance and can phosphorylate AMPK independently of AMP/ATP ratios^28^. As hypothalamic astrocytes and SCN circuits exhibit time-of-day-dependent Ca^2+^ dynamics^12–17^, we asked whether astrocytic Ca^2+^ signalling contributes to AMPK regulation in vivo using *IP_3_R2* knockout (*IP_3_R2KO*) mice, which lack astrocytic intracellular Ca^2+^ release^57,58^. In these mice, phosphorylation of AMPKα and its downstream target ACCα failed to exhibit the normal day-night variation observed in controls **(Fig. 6e and Supplementary Fig. 8c)**, indicating that astrocytic Ca^2+^ signalling contributes to the temporal regulation of this pathway. Consistent with this, phosphoproteomics revealed higher CaMKK2 phosphorylation at Ser495 during the night in control mice, whereas *γ2cKO* mice displayed elevated Ser495 phosphorylation already at ZT6, flattening its normal temporal profile **(Supplementary Fig. 8d)**. This effect was specific to CaMKK2, as no comparable day-night variation or genotype-dependent change was observed for CaMKK1 (Ser458).

Taken together, these findings support a model in which astrocytic Ca^2+^ signalling contributes to the temporal regulation of hypothalamic AMPK activity and fasting-responsive phosphosignalling networks, with CaMKK2 emerging as a candidate mediator of this effect **(Fig. 6f)**.

## Discussion

Our data show that hypothalamic astrocytic AMPK signalling displays time-of-day variation under constant darkness and in the absence of feeding cues, is modulated by nutrient availability and astrocytic Ca2+ dynamics, and contributes to the temporal organisation of the hypothalamic phosphoproteome. Disruption of astroglial AMPK signalling alters SCN-dependent circadian locomotor behaviour, PER2 abundance and phosphorylation, and is associated with reduced VIP expression. In addition, manipulations of astrocytic AMPK and PER2 in VMH astrocytes impair food-anticipatory activity. Together, these findings identify astrocytic AMPK signalling as a temporally regulated pathway linking metabolic state to hypothalamic timing and systemic homeostasis.

Across the experimental models examined, including gain-and loss-of-function manipulations of astroglial AMPK as well as pharmacological activation of AMPK in hypothalamic astrocytes, we observed coordinated changes in PER2 abundance, PER2 phosphorylation and AMPK signalling. *In vivo*, these alterations were accompanied by changes in free-running period, consistent with previous work showing that phosphorylation of specific PER2 residues can influence PER2 stability and circadian timing^11,12,35,59^. Although our data do not establish whether AMPK regulates PER2 directly or indirectly through CK1-dependent pathways^35^, they identify astroglial AMPK signalling as a regulator of PER2 temporal dynamics.

Classically viewed as an energy sensor regulated by AMP and ADP ratios^60–62^, AMPK in our study exhibited rhythmic regulation under constant darkness and in the absence of feeding cues, while remaining responsive to glucose availability. AMPK can be engaged by both metabolic and neuroendocrine signals: energy deficit activates the canonical AMP/ADP-sensitive pathway, whereas hormonal or neurotransmitter inputs can recruit Ca^2+^-dependent activation via CaMKK2^24–26,28^, consistent with our observations. Our findings demonstrate that astrocytic Ca²⁺ dynamics contribute to the temporal regulation of AMPK, with CaMKK2 providing a plausible link between astrocytic Ca^2+^ signalling and hypothalamic AMPK activation. Together these observations raise the possibility that astroglial AMPK signalling functions as an integrative node of circadian and metabolic inputs. Consistent with this interpretation, the phosphoproteomic alterations observed in *γ2cKO* hypothalami, involving pathways linked to circadian entrainment, insulin/PI3K–mTOR signalling and neurotransmission, together with the behavioural abnormalities and reduced SCN VIP expression, identify astroglial AMPK signalling as a regulator of hypothalamic phosphosignalling networks coordinating circadian and energy homeostasis.

The VMH integrates feeding-fasting state with circadian control^49–52^, and lesion studies have shown that VMH ablation suppresses FAA^53,63,64^. Our data extend this framework by identifying VMH astrocytic AMPK and PER2 in FAA. The altered postprandial AMPK phosphorylation profile observed in *γ2cKO* mice, together with the behavioural phenotypes of *γ2cKO* and *α1cKO* animals, indicates that appropriate temporal regulation of hypothalamic AMPK signalling is required for normal FAA expression, although the mechanistic relationships among VMH astrocytes, AMPKγ2 and PER2 remain unresolved.

Previous work has highlighted ghrelin, ketone bodies and insulin as candidate systemic cues for food-anticipatory rhythms, but current evidence does not assign a necessary or sufficient role to any single signal in driving FAA. This view is consistent with observations showing that insulin can synchronise astrocyte clock output and modulate food-entrainable behaviour^9^, whereas ghrelin and other metabolic hormones can activate hypothalamic AMPK and influence food-related motivation^2,65,66^. However, FAA can persist for several days under total food deprivation, even as circulating ghrelin or ketone levels lose their circadian cycling, indicating that hormone-dependent inputs are not strictly required for the maintenance of anticipatory rhythms. Moreover, systemic insulin sensitivity and hypothalamic insulin signalling exhibit circadian variation, and hypothalamic AMPK activity fluctuates across the day despite largely stable brain ATP levels. Together, these observations support a model in which multiple metabolic and neuronal signals converge on astrocytic AMPK-dependent pathways, with no single cue fully accounting for the circadian properties of food-anticipatory behaviour. Within this framework, our data point to nutrient availability and astrocytic Ca^2+^ dynamics as key modulators of hypothalamic AMPK activity in the context of scheduled feeding and fasting, while not excluding additional contributions from systemic hormones such as insulin or ghrelin. Thus, rather than a single upstream cue, astrocytic AMPK is likely to integrate multiple temporally patterned metabolic and neuronal inputs to support the timing mechanisms underlying FAA.

Taken together, our findings identify astrocytic AMPK signalling as a mechanism coupling metabolic state to circadian regulation through nutrient-sensitive and Ca²⁺-dependent pathways. Our data further suggest that temporally regulated astroglial AMPK signalling contributes to adaptive behavioural responses to scheduled feeding and may have broader implications for circadian-metabolic health, including sex-dependent vulnerability to shift work-associated metabolic disease^9,15,16,19,45,48^ and the efficacy of time-restricted feeding interventions^67^, where nutrient-responsive astroglial pathways may contribute to the cellular mechanisms linking feeding time to circadian and metabolic physiology.

## Methods

### Animals

All mice were bred and maintained at the animal facilities of the University of Santiago de Compostela in strict accordance with Spanish and European Union legislation on animal care. All procedures were approved by the University of Santiago de Compostela Ethics Committee, in compliance with Directive 2010/63/EU (Projects IDs 15012/2020/010 and 15012/2021/011). Mice were housed under standard conditions (12-h light/dark cycle, ZT0 = lights on at 08:00), in a temperature-and humidity-controlled environment, with *ad libitum* access to standard chow and water.

To generate conditional knockout (*cKO*) mice, *AMPKγ2^f/f^*(B6N.Cg-*Prkag2* tm1.1Rto/J, The Jackson Laboratory, #030979) and *AMPKα1^f/f^*(*Prkaa1*tm1.1Sjm/J, The Jackson Laboratory, #014141) lines were crossed with the *Glast-CreERT2* mice^30^. In some experiments, these mice were further crossed with Rosa26-tdTomato reporter mice (The Jackson Laboratory, #007914). *γ2cKO*, *α1cKO*, and littermate controls (*AMPKγ2^f/f^* or *AMPKα1^f/f^*) aged 12-16 weeks received oral gavage of tamoxifen (5 mg/day) dissolved in corn oil for two consecutive days to induce recombination^9,15,16,19,30,68^. Experiments were performed following a two-month tamoxifen washout period, according to the timelines specified for each assay. Adult male C57BL/6J mice (12-16 weeks old) were obtained from the institutional facility.

Hypothalamic samples from *IP_3_R2* knockout mice were obtained by João Filipe Oliveira from mice originally shared by Prof. Alfonso Araque (Minneapolis, MN, USA), under an agreement with Prof. Ju Chen (University of California, San Diego, CA, USA).

Primary hypothalamic astrocyte cultures were prepared from postnatal day 1-3 Sprague Dawley rats (Charles River Laboratories, Spain) of both sexes^17,18,69–73^. Pups were housed with their dams under standard conditions with *ad libitum* access to food and water.

### Stereotaxic microinjection of adeno-associated virus

To selectively manipulate gene expression in ventromedial hypothalamic (VMH) astrocytes, adult mice were subjected to stereotaxic surgery for bilateral injection of adeno-associated viruses (AAVs)^9,19^. Surgeries were performed under ketamine/xylazine anesthesia (15 mg/kg and 3 mg/kg, i.p.), and postoperative analgesia was provided with meloxicam (5 mg/kg) and buprenorphine (0.05 mg/kg). Mice were placed in a heated recovery cage and monitored until full recovery. Stereotaxic coordinates for VMH targeting were-1.7 mm posterior and ±0.5 mm lateral to bregma, and-5.5 mm ventral from the dura. For SCN targeting, coordinates were-0.5 mm posterior and ±0.3 mm lateral to bregma, and-6.0 mm ventral from the dura. Injections were performed bilaterally using a 33-gauge needle connected to a Hamilton syringe, delivering 2 × 10⁹ viral genome particles per side. To prevent backflow, the needle was left in place for 10 min before slow withdrawal. The incision was then closed with surgical sutures. All experiments were performed at least 3 weeks after injection to allow sufficient time for transgene expression or recombination.

To induce conditional deletion of AMPKγ2 in astrocytes, AAV2/5 vectors expressing Cre recombinase or EGFP under the human GFAP promoter (AAV-hGFAP-Cre and AAV-hGFAP-EGFP) were injected into the VMH or bilaterally into the SCN of *AMPKγ2^fl/fl^* mice^9,19^. To achieve astrocyte-specific cell ablation, AAV2/5 expressing a Cre-dependent truncated caspase-3 construct (AAV-flex-taCasp3-TEVp; Addgene #45580) was injected into the VMH of Glast-CreERT2-tdTomato mice. This construct induces Cre-dependent, cell-autonomous apoptosis, thereby minimizing toxicity to neighbouring non-Cre-expressing cells^74^. To knock down *Per2* in astrocytes, an AAV5 vector expressing a miR30-based short hairpin RNA (shRNA) targeting *Per2* under the GFAP (short) promoter was injected into the VMH. The shRNA sequence used was AGTGGACAGCCTTTCGATTATTTAGTGAAGCCACAGATGTAAATAATCGAAAGGCTGTCCACC.

### Circadian locomotor activity

Circadian activity rhythms were assessed in 4-to 5-month-old *γ2cKO*, *α1cKO*, and control mice using running wheel assays. Mice were single-housed in ventilated cages equipped with low-profile wireless running wheels (ENV-044; Med Associates Inc.). Males and females were housed separately on independent ventilated racks to prevent sex-related olfactory or social interference. All cages were maintained under uniform lighting conditions in the same room to ensure consistent light exposure across groups (14-16, 18, 19, 38). Animals were acclimated to the wheels for 3 days under a 12 h light/12 h dark (LD) cycle, followed by 7-10 days of LD recording. Mice were then transferred to fully blacked-out cages (Tecniplast) and maintained in constant darkness (DD) for 3-4 weeks^9,15,16,19,45^.

Wheel-running activity was recorded in 5-min bins using Wheel Manager software (SOF-860; Med Associates Inc.) and analyzed with ActogramJ. LD activity was plotted using zeitgeber time (ZT), with ZT0 defined as lights on. For DD conditions, data were plotted using circadian time (CT), adjusted according to each animal’s endogenous period. Mean activity profiles were generated by averaging 5-min-binned activity over 7-10 consecutive stable days. Free-running period and rhythm amplitude were estimated with Lomb-Scargle periodograms. Female data were collected across multiple estrous cycles to account for intra-individual variability^9,15,16,19,45^

### Food restriction experiments

Food-entrainable circadian rhythms were assessed in control, *γ2cKO*, and *α1cKO* mice, 14 weeks after tamoxifen-induced recombination, as well as mice subjected to stereotaxic VMH injections of AAV-Cre (*AMPKγ2^f/f^*), AAV-flex-taCasp3-TEVp (*Glast-CreERT2*), or shRNA targeting *Per2* (AAV[miR30]-GFAP-EGFP) and respective controls. All mice were single-housed in cages equipped with running wheels (ENV-044; Med Associates Inc.) under 12 h:12 h LD conditions. Water was available *ad libitum* throughout restricted feeding. Prior to restricted feeding, baseline wheel-running activity was recorded for 7 days under *ad libitum* feeding. Body weight and food intake were measured at zeitgeber time 4 (ZT4). Mice were then subjected to a 10-day RF schedule: during the first 6 days, food access was limited to ZT4-ZT10; for the remaining 4 days, it was further restricted to ZT4-ZT8^9,19,45^. Standard chow (Envigo) was provided during these windows. Food consumption was monitored daily by weighing pellets at the beginning (ZT4) and end (ZT10 or ZT8) of each feeding period, with values corrected for spillage. Locomotor activity was recorded in 5-min bins using Wheel Manager software (SOF-860; Med Associates Inc.) and analyzed with ActogramJ^9,19,45^.

### Metabolic and physiological assessments

Body weight and food intake were recorded weekly. Whole-body composition was determined by nuclear magnetic resonance (NMR) using an EchoMRI analyzer (EchoMRI, Houston, TX)^9,19,45,49,75,76^. Skin temperature over the interscapular brown adipose tissue (BAT) was assessed via infrared thermography (FLIR B335 camera; FLIR Systems) and analyzed with FLIR Tools software. For each animal, three dorsal images were acquired, and mean BAT surface temperature was calculated^9,19,45,49,75,76^. Measurements were performed at a fixed zeitgeber time (ZT8) to control for circadian variation. Rectal temperature was measured with a digital thermistor probe immediately before thermal imaging. Energy expenditure, respiratory exchange ratio (RER), and locomotor activity were measured using an open-circuit indirect calorimetry system (LabMaster; TSE Systems). Mice were acclimated to metabolic cages for 24 h, before data acquisition over the subsequent 48-72 _h9,19,45,49,75,76._

### Glucose and insulin tolerance test

For glucose tolerance tests (GTT), mice were fasted for 6 h and injected intraperitoneally with D-glucose (2 g/kg; Sigma-Aldrich, G8270). For insulin tolerance tests (ITT), human insulin (Actrapid; Novo Nordisk) was administered at 0.75 IU/kg in males and 0.35 IU/kg in females. Blood glucose levels were measured at designated time points using a handheld glucometer^9,15,19,45,49,75,76^.

### Western blotting

Protein expression analyses were performed on tissue samples from the hypothalamus, mediobasal hypothalamus (MBH), brown adipose tissue (BAT), and on primary hypothalamic astrocytes derived from neonatal rats. Samples were homogenized in ice-cold lysis buffer containing 50 mM Tris-HCl, 10 mM EGTA, 1 mM EDTA, 1.6% Triton X-100, 1 mM sodium orthovanadate, 50 mM sodium fluoride, 10 mM sodium pyrophosphate, and 250 mM sucrose (pH 7.5). Protease inhibitor cocktail (Roche Diagnostics) was added fresh prior to homogenization^9,15–17,19,45,49,75,76^. Protein concentration was determined using the Bradford assay (Bio-Rad). Equal amounts of protein were separated by SDS-PAGE and transferred to nitrocellulose membranes (Protran, Schleicher & Schuell). Membranes were blocked in 5% non-fat dry milk in TBS-T (20 mM Tris-HCl, 150 mM NaCl, 0.1% Tween-20) for 1 h at room temperature. Membranes were incubated with primary antibodies against: PER2 (1:1000, Invitrogen PA5-89045), phospho-PER2 (Ser662) (1:1000, Invitrogen PA538901), CRY1 (1:1000, Invitrogen PA5-89349), UCP1 (1:1000, Abcam ab10983), phospho-AMPKα (1:1000, Cell Signaling #2535S), total AMPKα (1:1000, Cell Signaling #2532), AMPKγ1 (1:1000, Thermo Fisher PA5-27471), VIP (1:500, Invitrogen PA5-78224), AVP (1:1000, Santa Cruz Biotechnology sc-390723), phospho-ACCα (1:1000, Cell Signaling #3661S), and total ACCα (1:1000, Cell Signaling #3662S). HRP-conjugated secondary antibodies (sc-2357 and sc-525408; Santa Cruz Biotechnology) were used at 1:2000. After detection, membranes were stripped (Thermo Fisher, #46430) and re-probed with loading control antibodies: GAPDH (1:5000, Santa Cruz Biotechnology sc-25778), β-actin (1:5000, Sigma-Aldrich A5316), or α-tubulin (1:5000, Sigma-Aldrich AB_330337), depending on tissue source. Immunoreactive bands were visualized using enhanced chemiluminescence (ECL; GE Healthcare) and exposed to autoradiographic film (Fujifilm). Band intensity was quantified by densitometry using ImageJ 1.33 (NIH). Signals were normalized to GAPDH (primary astrocytes), β-actin (hypothalamus), or α-tubulin (BAT). Representative images are shown for each target protein. All bands shown originate from the same gel, with digital repositioning applied when necessary for clarity^9,15–17,19,45,49,75,76^.

### Immunofluorescence

Mice were deeply anesthetized with ketamine/xylazine (150 mg/kg and 10 mg/kg, i.p.) and transcardially perfused with ice-cold phosphate-buffered saline (PBS), followed by 4% paraformaldehyde in PBS. Brains were post-fixed overnight in 4% PFA at 4 °C, then coronally sectioned at 30 μm using a cryostat (Leica Microsystems). Free-floating sections were permeabilized with 0.3% Triton X-100 in PBS, blocked with 10% goat serum for 1 h at room temperature, and incubated overnight at 4 °C with rabbit anti-VIP (1:500, Invitrogen PA5-78224) and phospho-ACCα (1:50, Cell Signaling #3661S). The following day, sections were washed and incubated for 2 h at room temperature with Alexa Fluor-conjugated secondary antibodies (Alexa Fluor 647 and Alexa Fluor 488). Slices were mounted using ProLong Gold Antifade Mountant (Thermo Fisher Scientific, P36934) and imaged with a Leica TCS SP5 inverted laser scanning confocal microscope equipped with 20× and 40× objectives. Quantification was performed using ImageJ (NIH). Regions of interest (ROIs), including the SCN, VMH and ARC, were delineated based on DAPI nuclear staining, and the average fluorescence intensity of the target signal was measured. For each animal, the mean value from multiple consecutive sections was used as a single biological replicate^9,15–19,45,68^.

### Immunohistochemistry

Brown adipose tissue samples were fixed in 10% neutral-buffered formalin for 24 h, dehydrated, and embedded in paraffin using standard procedures. Serial sections (3 μm) were obtained with a rotary microtome. For immunohistochemical detection of UCP1, sections were incubated with a rabbit anti-UCP1 primary antibody (1:2000; Abcam ab10983) followed by appropriate secondary antibodies. UCP1-positive cells were visualized and quantified using ImageJ, based on thresholded staining _intensity9,19,49,75,76._

### Primary hypothalamic astrocyte cultures

Primary monolayer cultures of astrocytes were established from the hypothalamus of neonatal (P1-P3) Sprague Dawley rats^9,16–18,69–73^. Neonatal rat pups were decapitated using sharp scissors. Astrocyte cultures were maintained at 37 °C in a humidified 5% CO₂ atmosphere for 1 week. After 1 week, cells were trypsinized and subcultured for experiments. These astrocyte-enriched cultures contained >96% astrocytes, as determined by immunofluorescence using a monoclonal anti-GFAP antibody (clone 6F2, Dako; 1:1000 dilution)^9,16–18,69–73^. Astrocytes were treated with AICAR (1 mM) and/or cycloheximide (CHX, 50 μg/mL) in medium containing either low (1000 mg/L) or high (4500 mg/L) glucose.

### Phosphoproteomic sample preparation

Hypothalamic tissues were homogenized using a FastPrep-24™ 5G instrument (MP Biomedicals) in lysis buffer containing 7 M urea, 2 M thiourea, 4% CHAPS, 200 mM DTT, and phosphatase inhibitor cocktail. Following sonication and centrifugation, proteins were precipitated with acetone overnight; this precipitation step was repeated three times. After the final precipitation, protein pellets were resuspended in lysis buffer lacking phosphatase inhibitors and quantified. Proteins were digested following a modified filter-aided sample preparation (FASP) protocol as described by Wiśniewski et al., with minor modifications^77^. Trypsin was added at a 1:20 enzyme-to-substrate ratio and incubated overnight at 37 °C. Resulting peptides were dried in a vacuum concentrator and resuspended in 0.1% formic acid. Phosphopeptides were enriched using the High-Select™ TiO₂ Phosphopeptide Enrichment Kit (Thermo Fisher Scientific) according to the manufacturer’s instructions (∼500 μg protein input per sample). Eluted phosphopeptides were directly subjected to LC-MS/MS analysis.

### Phosphopeptide mass spectrometry and data analysis

Phosphopeptide samples (∼20% of enriched material) were loaded onto an Evosep One system (Evosep) coupled online to a timsTOF Pro mass spectrometer using PASEF acquisition (Bruker Daltonics). Separation was performed using the 30 samples-per-day Evosep protocol. Raw data were processed using FragPipe against the *Mus musculus* UniProt/Swiss-Prot database, with precursor and fragment mass tolerances of 20 ppm and 0.05 Da, respectively. Carbamidomethylation of cysteine was set as a fixed modification, whereas methionine oxidation and phosphorylation of serine, threonine, and tyrosine were specified as variable modifications. Peptide identifications were filtered at a 1% false discovery rate (FDR). Quantitative values from FragPipe were log₂-transformed and analyzed in Perseus^78^. Phosphopeptides present in at least 70% of samples in any experimental group were retained for further analysis. Missing values were imputed based on a normal distribution of the lowest 10% of signal intensities within each sample.

### In silico analysis

*In silico* analyses were performed on hypothalamic phosphoproteomic datasets obtained from mice subjected to 24 h of food deprivation. Animals were sacrificed at four circadian time points (ZT0, ZT6, ZT12, and ZT18) to capture temporal phosphorylation dynamics across the light-dark cycle. Additionally, phosphoproteomic profiles were generated from the hypothalami of control and *γ2cKO* mice, collected at ZT6 and ZT18, to assess genotype-specific effects on circadian phosphorylation. To identify circadian rhythmic phosphorylation events, data from starved wild-type mice were analyzed using BIO_CYCLE^79,80^, which detects rhythmicity based on periodic regression models. Phosphopeptides were considered rhythmic if they exhibited significant fits (q < 0.05, Benjamini-Hochberg). When multiple peptides mapped to the same protein, the one with the highest rhythmic amplitude was retained for downstream analysis. Rhythmic phosphopeptides were grouped by peak phase and clustered based on z-scored intensity profiles across time points ^9^. The number of temporal clusters was determined by minimizing the Bayesian information criterion (BIC).

Functional enrichment analysis was performed using Gene Ontology (GO) and Kyoto Encyclopedia of Genes and Genomes (KEGG) pathway annotations via the DAVID Bioinformatics Resource, applying default parameters for GO Biological Process (BP_FAT) terms to prioritize specific over broad categories^9^. Kinase-substrate enrichment analysis was conducted using KEA3 (Kinase Enrichment Analysis 3), which predicts upstream kinase activity based on the overrepresentation of known kinase-substrate relationships in the input phosphopeptide set. Default settings were used throughout.

## Statistical analysis

Data are presented as mean ± s.e.m. and were analyzed using GraphPad Prism 10 (GraphPad Software, San Diego, CA, USA). Exact n values, statistical tests, and measures of variability are reported in the figure legends. Statistical significance was assessed using multiple approaches depending on the experimental design. For paired or unpaired comparisons, two-tailed Student’s t-tests were used as appropriate. Two-way repeated-measures ANOVA with Bonferroni post hoc correction was used for time-course and repeated-measures data. All datasets were tested for normality and homogeneity of variance prior to analysis. Rhythmicity in gene or protein expression was evaluated using Cosinor analysis, and circadian parameters (period, phase, mesor, amplitude) were extracted accordingly^9,15–19,45,68^.

In food-restriction experiments, food anticipatory activity (FAA) was quantified as the duration (in hours) of wheel-running activity during the pre-feeding period. The FAA ratio was calculated as the fold change in activity during the 4 h preceding food availability relative to the rest of the day. Similarly, the nocturnal activity ratio was defined as the fold change in activity during the dark phase relative to total daily activity^9,19,45^.

## Acknowledgements

We thank Dr. M. Götz (Physiological Genomics, Biomedical Center, Ludwig-Maximilians-University Munich, Germany) for providing the *Glast-CreERT2* mouse line. We also thank Prof. Michael Hastings and Dr. Andrew Patton (MRC Laboratory of Molecular Biology, Cambridge, UK), and Prof. Rubén Nogueiras (CiMUS, University of Santiago de Compostela, Spain) for scientific discussions. We thank the technical staff at CiMUS for support, and the USC Animal Facility (CEBEGA) for assistance with animal care and experimentation. Some figures were created using BioRender.com (2025).

## Author contributions

O.B.-M. and M.L. conceptualized the study. O.B.-M. designed the experimental framework and supervised the overall project. M.L. generated and provided the *γ2cKO* and *α1cKO* mouse lines. M.L.-M. and A.G.-V. performed most of the experiments. O.B.-M., M.L.-M. and A.G.-V. analysed and interpreted the data. M.S.-L. carried out the metabolic characterization of the mouse models. A.M.I.-A., V.P.-G., H.L.-G., H.C.M., P.M.-P., N.R.V.D., P.F.-S. and A.M.E.-S. provided technical support and assisted in sample collection and western blot analyses. J.F.O. and D.S.A. contributed hypothalamic samples from IP3R2 knockout mice. F.E. and M.Az. conducted the phosphoproteomic experiments and bioinformatic analyses. O.B.-M. wrote the manuscript with input from all authors.

## Data and materials availability

All data supporting the findings of this study are available from the corresponding authors upon reasonable request.

