## Supplementary Figures for "Astrocytic AMPK couples metabolic signals to hypothalamic circadian timing"

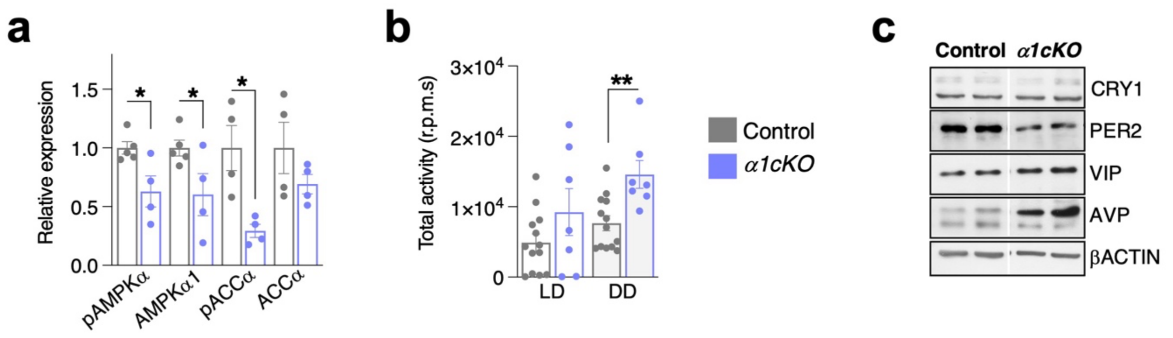


**Supplementary Fig. 1. Astrocytic AMPKα1 deletion is associated with altered hypothalamic AMPK signalling and total locomotor activity**. **a,** Quantification of pAMPKα, AMPKα1, pACCα and ACCα protein levels in hypothalamic lysates from control and *α1cKO* mice collected at ZT8 (n = 4-5 biologically independent animals per group). **b,** Total locomotor activity under LD and DD conditions (control n = 13, *α1cKO* n = 7). **c**, Representative western blots of CRY1, PER2, VIP and AVP in hypothalamic lysates from control and *α1cKO* mice collected at ZT8. Data are presented as mean ± s.e.m. Statistical significance was determined using unpaired two-tailed t-tests. *P < 0.05, **P < 0.01.

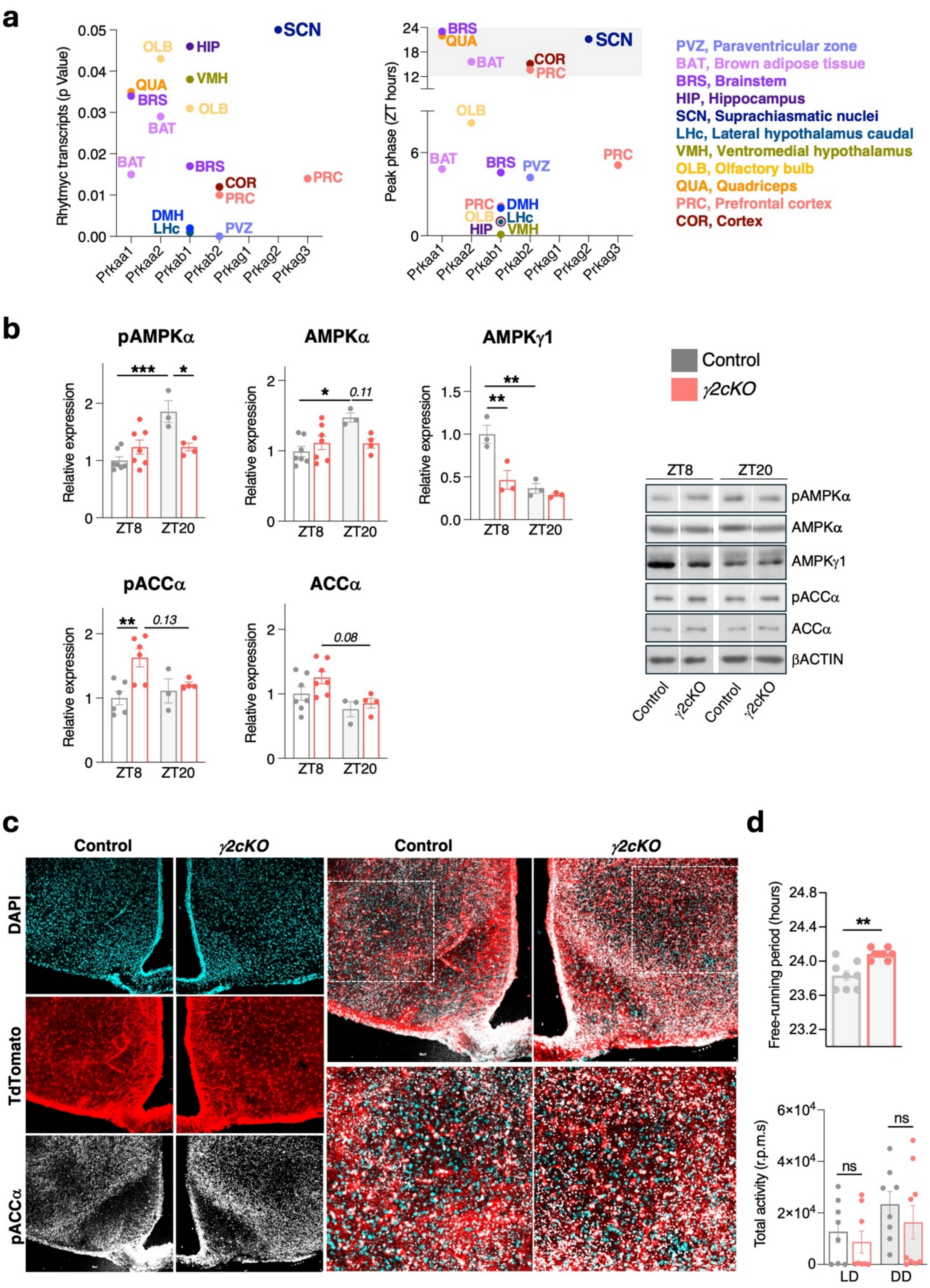


**Supplementary Fig. 2. Astrocytic AMPKγ2 deletion is associated with altered hypothalamic AMPK signalling and circadian behaviour. a,** Reanalysis of published transcriptomic data from CBA/CaJ mice showing rhythmic expression and peak phase of AMPK subunit genes across multiple brain regions. *Prkag2* displayed prominent rhythmic expression in the SCN with peak levels during the night. Left: significance of rhythmicity (JTK_CYCLE *q*-value); right: peak phase distribution of AMPK subunit genes across brain regions. **b,** Western blot analysis of hypothalamic tissue from control and *γ2cKO* mice collected at ZT8 and ZT20. Quantification of pAMPKα, AMPKα, AMPKγ1, pACCα, and ACCα protein levels (left) and representative immunoblots (right) (n = 3-7). **c,** Representative immunofluorescence images showing pACCα in the VMH and arcuate nucleus (ARC) at ZT0. Tomato fluorescence marks recombined astrocytes in *γ2cKO; tdTomato* reporter mice. **d,** Quantification of free-running period (upper panel) and total locomotor activity (lower panel) under LD and DD conditions (n = 8). Data are presented as mean ± s.e.m. Statistical significance was determined using two-way ANOVA (b) and unpaired two-tailed t-tests (d). ns, not significant; *P < 0.05, **P < 0.01, ***P < 0.001.

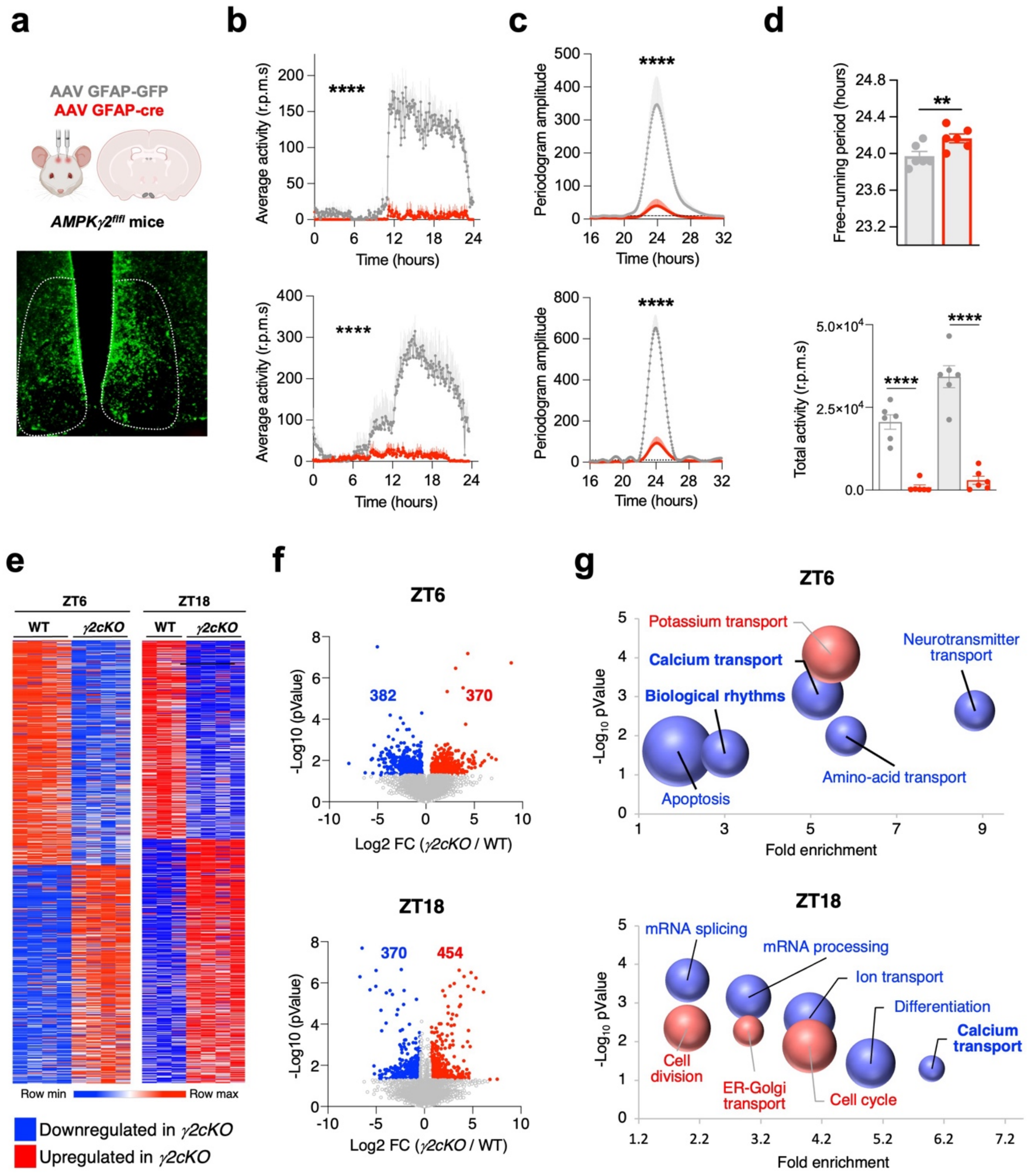


**Supplementary Fig. 3. Astrocytic AMPKγ2 deletion disrupts circadian locomotor rhythms and alters the hypothalamic phosphoproteome.** **a**, Schematic of SCN-targeted astrocytic AMPKγ2 deletion using AAV-GFAP-Cre in *AMPKγ2*$^{fl/fl}$ mice and representative fluorescence image showing viral targeting within the SCN. Average locomotor activity profiles **(b)** and corresponding Lomb-Scargle periodogram analysis **(c)** of control and SCN-*γ2KO* mice under LD and DD conditions (top and bottom panels, respectively) (n = 6). **d,** Quantification of free-running period (upper panel) and total locomotor activity (lower panel) in control and *SCN-γ2KO* mice (n = 6). **e,** Heatmaps showing phosphopeptide abundance in control and *γ2cKO* hypothalami collected at ZT6 and ZT18 (control n = 3; *γ2cKO* n = 4). **f,** Volcano plots showing differentially phosphorylated sites between genotypes at ZT6 (top) and ZT18 (bottom). **g,** Gene Ontology biological process enrichment analysis of phosphoproteins displaying genotype-dependent differences at ZT6 and ZT18. Bubble size reflects $-\log_{10}$(P value), whereas position indicates fold enrichment. Data are presented as mean ± s.e.m. Statistical significance was determined using two-way ANOVA (b, c) and unpaired two-tailed t-tests (d). $^{**}P < 0.01$, $^{****}P < 0.0001$.

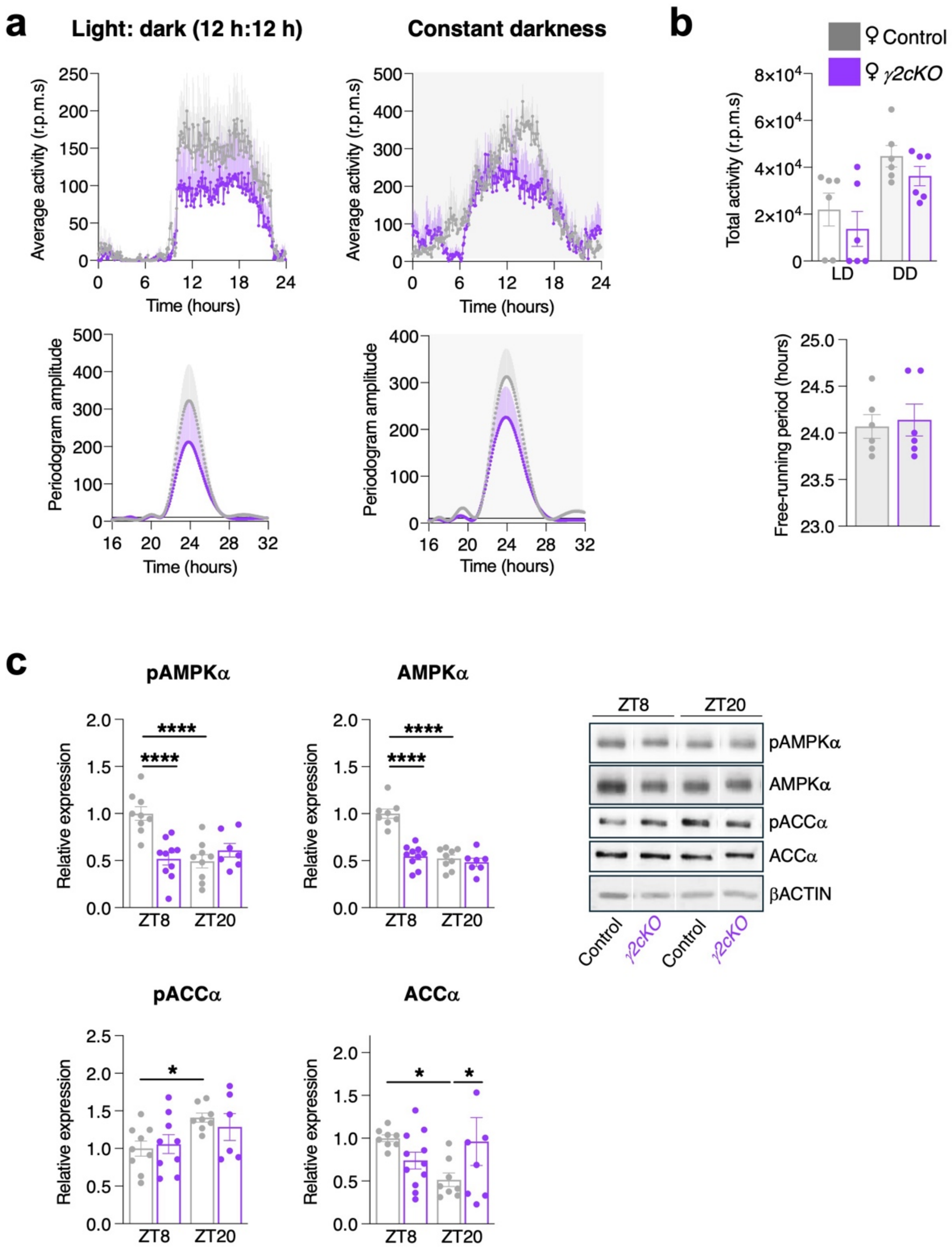


**Supplementary Fig. 4. Female *γ2cKO* mice exhibit altered hypothalamic AMPK signalling with largely preserved circadian behaviour. a,** Average locomotor activity profiles of control and *γ2cKO* female mice under LD and DD conditions (upper panels). Lomb-Scargle periodogram analysis of locomotor activity rhythms under LD and DD conditions (lower panels) (n = 6). **b,** Total activity counts under LD and DD conditions (upper panel) and free-running period (lower panel) in control and *γ2cKO* female mice (n = 6). **c,** Quantification of pAMPKα, total AMPKα, pACCα and total ACCα protein abundance in hypothalamic lysates collected from control and *γ2cKO* female mice at ZT8 and ZT20 (left), with representative immunoblots shown on the right (n = 6-11). Data are presented as mean ± s.e.m. Statistical significance was determined using two-way ANOVA and unpaired two-tailed t-tests. *P < 0.05, ****P < 0.0001.

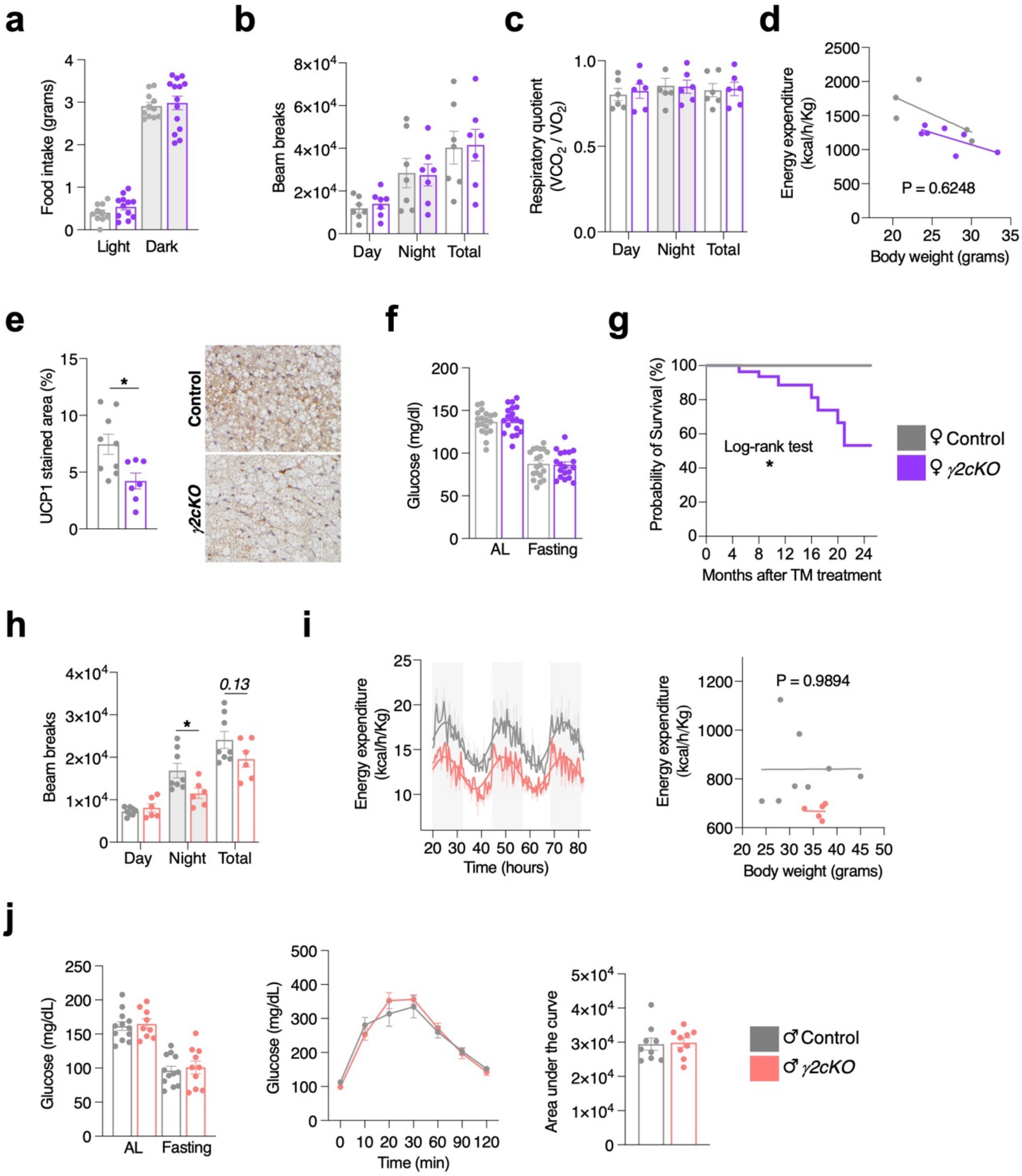


**Supplementary Fig. 5. Additional metabolic, locomotor and survival phenotypes in *γ2cKO* mice.** **a**, Food intake during light and dark phases in control and *γ2cKO* females (n = 11-14). **b, h,** Locomotor activity (beam breaks) during the light and dark phases and total counts in females (b, n = 7) and males (h, n = 6-8), respectively. **c**, Respiratory quotient ($VCO_2/VO_2$) during the light and dark phases and mean values in females (n = 5-6). **d, i,** Correlation between energy expenditure and body weight in females (d, n = 5-7) and males (i, n = 5-8). Representative 72 h energy expenditure profiles are shown in **i** (left). **e**, Quantification of UCP1-positive area in BAT and representative immunohistochemical images from control and *γ2cKO* females (n = 7-9). **f**, *Ad libitum* (AL)-fed and fasting blood glucose levels in females (n = 19). **g**, Survival analysis of control and *γ2cKO* females after tamoxifen (TM) treatment (n = 101). **j**, AL-fed and fasting blood glucose levels (left), glucose tolerance test (middle) and corresponding area under the curve (right) in males (n = 9-12). Data were collected 20 weeks after TM induction. Data are presented as mean ± s.e.m. Statistical significance was determined using two-way ANOVA, unpaired two-tailed t-tests and the log-rank test. *$P < 0.05$.

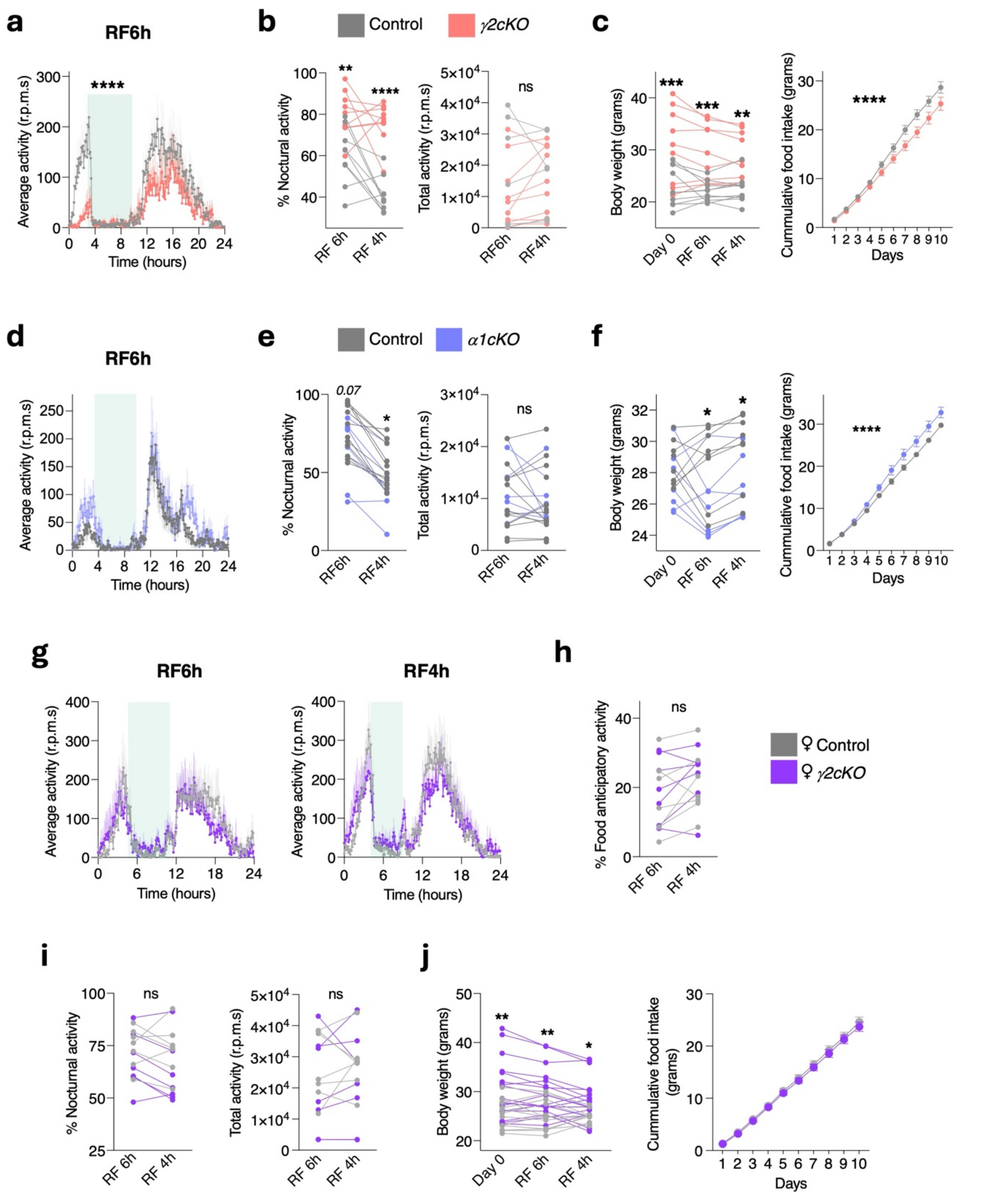


**Supplementary Fig. 6. Astrocytic AMPK signalling contributes to behavioural and metabolic adaptation during restricted feeding (RF). a,** Average locomotor activity profiles of control and *γ2cKO* male mice during RF with a 6 h food access window (RF6h) (n = 7). **b,** Quantification of nocturnal activity (left) and total locomotor activity (right) in control and *γ2cKO* male mice after adaptation to RF6h and RF4h conditions (n = 7). **c,** Body weight (left) and cumulative food intake (right) in control and *γ2cKO* male mice during restricted feeding (n = 7-9). **d-f,** Same as in **a-c**, respectively, for control and *α1cKO* male mice (n = 6-10). **g,** Average locomotor activity profiles of control and *γ2cKO* female mice under RF6h and RF4h conditions (n = 6-7). **h,** Quantification of FAA in control and *γ2cKO* female mice after adaptation to RF6h and RF4h conditions (n = 6-7). **i,** Quantification of nocturnal activity (left) and total locomotor activity (right) in control and *γ2cKO* female mice under RF conditions (n = 6-7). **j,** Body weight (left) and cumulative food intake (right) in control and *γ2cKO* female mice during RF (n = 6-7). Data are presented as mean ± s.e.m. Statistical significance was determined using two-way ANOVA. *P < 0.05, **P < 0.01, ***P < 0.001, ****P < 0.0001; ns, not significant.

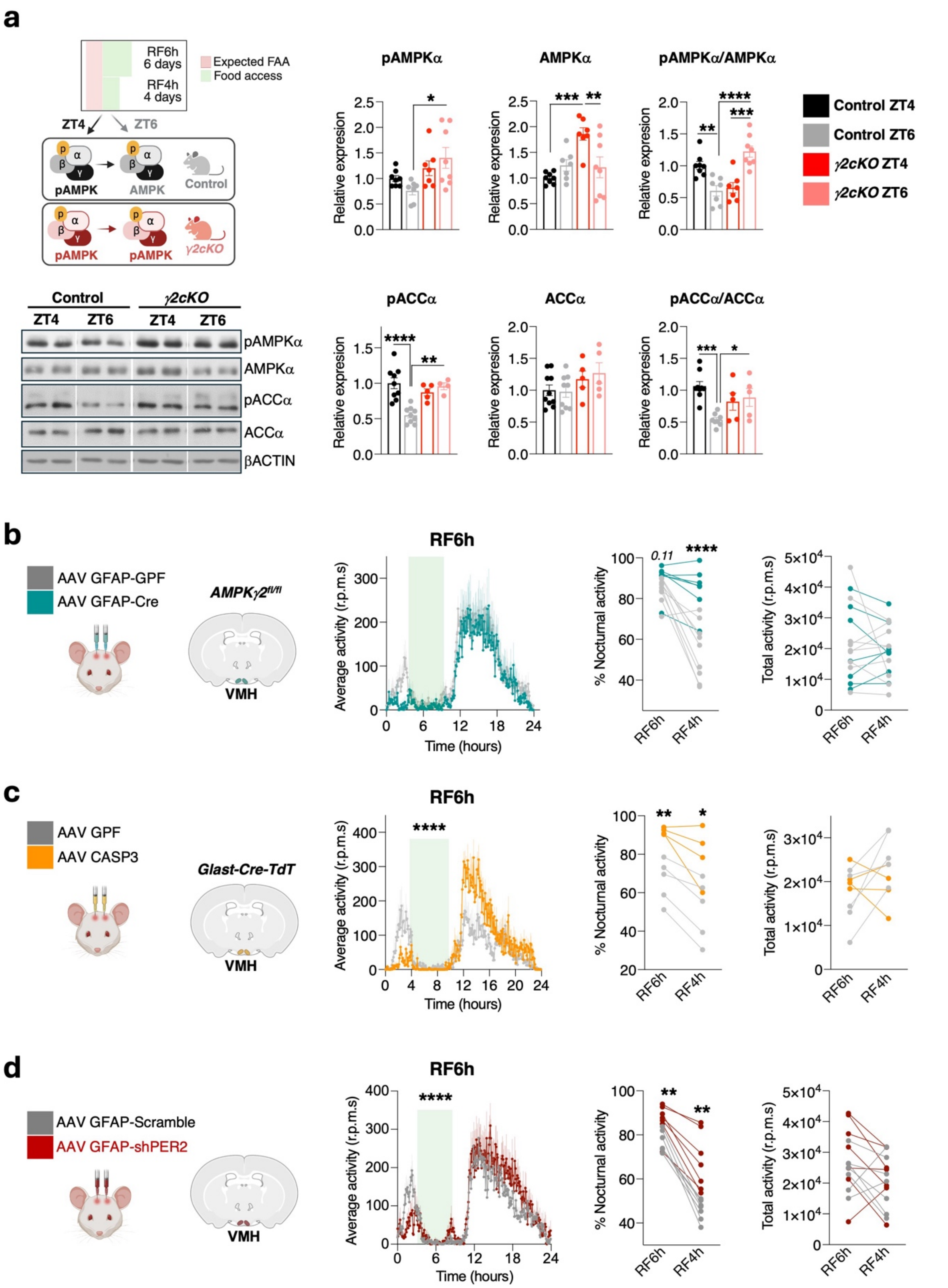


**Supplementary Fig. 7. VMH astrocytic AMPK and PER2 signalling contribute to FAA. a,** Schematic of the RF paradigm and experimental design for analysis of hypothalamic AMPK signalling at ZT4 and ZT6 (left). Representative immunoblots of hypothalamic pAMPKα, AMPKα, pACCα and ACCα protein levels in control and *γ2cKO* male mice collected at ZT4 and ZT6 (bottom). Quantification of pAMPKα, AMPKα, pAMPKα/AMPKα ratio, pACCα, ACCα and pACCα/ACCα ratio is shown on the right (n = 4-10). **b,** Schematic of VMH-targeted astrocytic AMPKγ2 deletion using AAV GFAP-Cre in *AMPKγ2$^{fl/fl}$* mice (left). Average locomotor activity profiles during RF6h (middle), and quantification of nocturnal activity and total locomotor activity (right) following VMH-specific AMPKγ2 deletion (n = 6-9). **c,** Schematic of VMH astrocyte ablation using AAV-flex-taCASP3-TEVp in *Glast-Cre-TdTomato* mice (left). Average locomotor activity profiles during RF6h (middle), and quantification of nocturnal activity and total locomotor activity (right) following VMH astrocyte ablation (n = 4-5). **d,** Schematic of astrocytic PER2 knockdown in the VMH using AAV GFAP-shPER2 (left). Average locomotor activity profiles during RF6h (middle), and quantification of nocturnal activity and total locomotor activity (right) following astrocytic PER2 knockdown (n = 6-7). Data are presented as mean ± s.e.m. Statistical significance was determined using two-way ANOVA. *P < 0.05, **P < 0.01, ***P < 0.001, ****P < 0.0001.

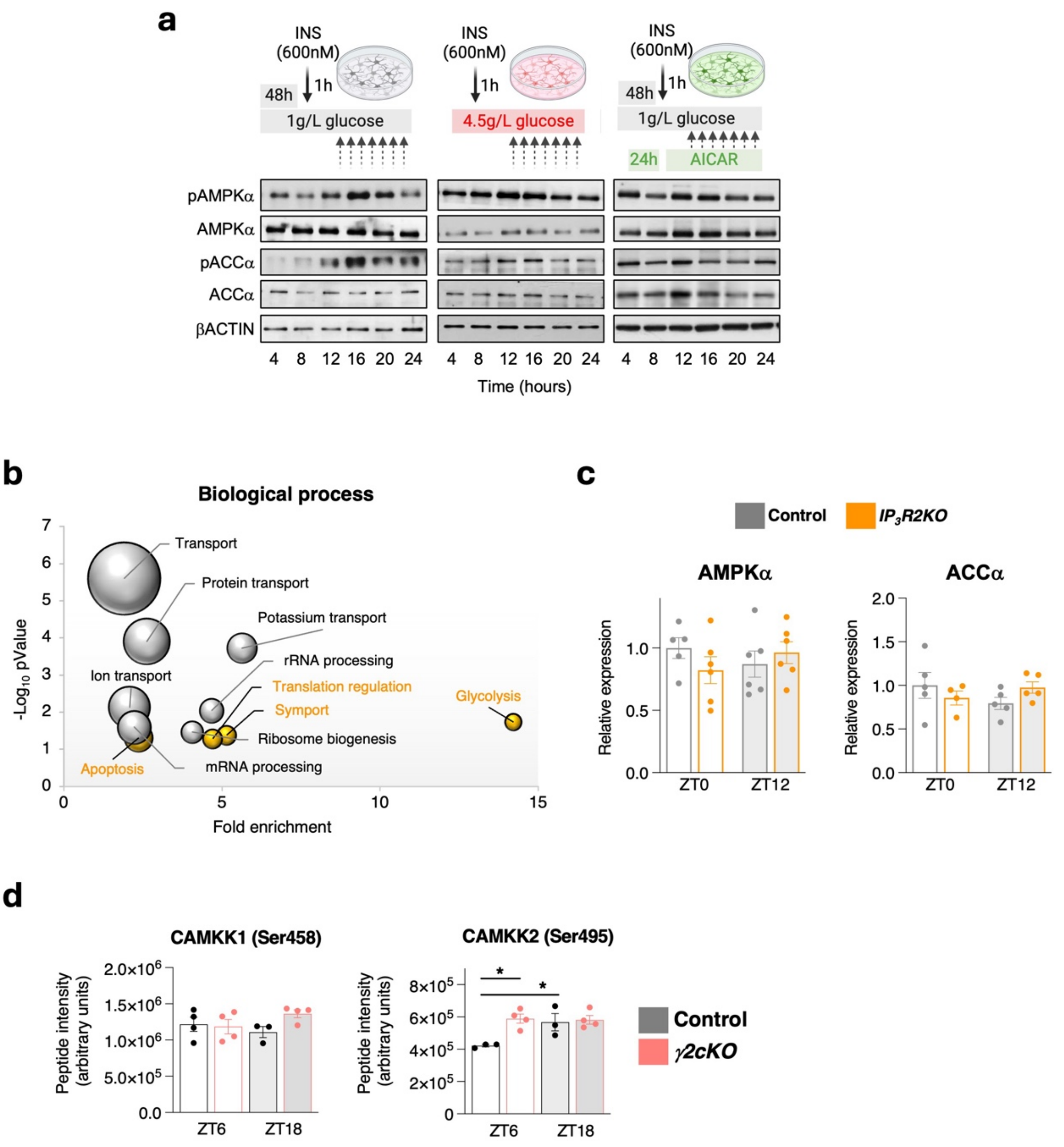


**Supplementary Fig. 8. Astrocytic intracellular $Ca^{2+}$ contributes to temporal hypothalamic AMPK signalling. a,** Representative immunoblots of pAMPKα, AMPKα, pACCα and ACCα in primary hypothalamic astrocytes synchronized with insulin (600 nM, 1 h) and maintained under low-glucose (1 g/L; left), high-glucose (4.5 g/L; middle) or AICAR-treated low-glucose conditions (right). Samples were collected at the indicated time points after synchronization. Representative of two independent experiments. **b,** Gene Ontology enrichment analysis of biological processes associated with rhythmic phosphoproteins in 24-h fasted hypothalami. Daytime-enriched terms are shown in yellow and night-time-enriched terms in grey. Bubble position indicates fold enrichment and statistical significance ($-\log_{10}$(P value)), whereas bubble size reflects the number of proteins assigned to each term. **c,** Quantification of total AMPKα and ACCα protein levels in hypothalamic extracts from control and *$IP_3R2KO$* mice collected at ZT0 and ZT12 under *ad libitum* feeding conditions (n = 4-6 per group). **d,** Phosphorylation levels of CaMKK1 (Ser458) and CaMKK2 (Ser495) in hypothalamic lysates from control and *γ2cKO* mice collected at ZT6 and ZT18 (n = 3-4 per group). Data are presented as mean ± s.e.m. Statistical significance was determined using two-way ANOVA. $^*P < 0.05$.
